# Mechanical loading is required for initiation of extracellular matrix deposition at the developing murine myotendinous junction

**DOI:** 10.1101/2022.06.28.497966

**Authors:** Sarah N. Lipp, Kathryn R. Jacobson, Haley A. Colling, Tyler G. Tuttle, Dalton T. Miles, Kaitlin P. McCreery, Sarah Calve

## Abstract

The myotendinous junction (MTJ) contributes to the generation of motion by connecting muscle to tendon. At the adult MTJ, a specialized extracellular matrix (ECM) is thought to contribute to the mechanical integrity of the muscle-tendon interface, but the factors that influence MTJ formation during mammalian development are unclear. Here, we combined 3D imaging and proteomics with murine models in which muscle contractility and patterning are disrupted to resolve morphological and compositional changes in the ECM during MTJ development. We found that MTJ-specific ECM deposition can be initiated via static loading due to growth; however, it required cyclic loading to develop a mature morphology. Furthermore, the MTJ can mature without the tendon terminating into cartilage. Based on these results, we describe a model wherein MTJ development depends on mechanical loading but not insertion into an enthesis.

## Introduction

To generate movement, force produced by skeletal muscle contraction is transmitted from tendon to bone. The interface between muscle and tendon, the myotendinous junction (MTJ), is organized to withstand repeated cycles of muscle-generated forces. The ends of myofibers at the MTJ terminate in a highly interdigitated interface with tendon, which is linked by specialized cell-cell and cell-extracellular matrix (ECM) interactions [1].

The ECM is a network of proteins and glycosaminoglycans that contributes to tissue formation and function by regulating mechanical stability, cell signaling, and growth factor availability [2]. Individual tissues have distinct biomechanical demands, which is reflected by differences in ECM composition and organization [2]. For example, type XXII collagen (COL22A1) and periostin (POSTN) are localized to the adult MTJ, but not found in the adjacent muscle and tendon [3]. Mutations in COL22A1 significantly correlate with muscle injury in humans, indicating the ECM is critical for maintaining adult MTJ functionality [4]. Correct assembly of the ECM is also important for MTJ development as demonstrated in studies where thrombospondin-4 (THBS4), COL22A1, and POSTN were knocked down in zebrafish [5–7]. Despite the essential role in facilitating motion, few studies have investigated the factors necessary for mammalian MTJ formation.

Various interactions are required for the correct assembly of each muscle-tendon-bone complex. One type of interaction is force transmission, which includes semi-static loading via the rapid, anisotropic skeletal growth that induces tension on the developing muscle-tendon unit [8, 9], and cyclic loading from muscle contraction [10]. Reciprocal signaling between different cell and tissue types also contributes to musculoskeletal assembly. Muscle patterning is influenced by connective tissue cells regulated by transcription factors such as *Osr1, Tbx3, Tbx5*, and *Tcf4* [11–14]. For example, mutations in *Tbx3* disrupt dorsal patterning, leading to an abnormal lateral triceps muscle origin and insertion [14]. In contrast, the initial patterning of tendons is thought to be regulated by signaling from the developing skeleton [10]. Subsequent maturation of both muscle and tendon requires muscle contraction, as indicated by studies using a murine model of muscular dysgenesis (*mdg*) in which a mutation in a calcium ion channel (CACNA1S) disrupts muscle excitation-contraction coupling [10, 15, 16]. Investigations in zebrafish and *Drosophila* demonstrated muscle contraction is also critical for MTJ maturation [17, 18]. However, the roles of mechanical loading and reciprocal interactions between muscle, tendon, and cartilage/bone during mammalian MTJ formation and ECM patterning remain unclear.

The objectives of this work were to (1) characterize the deposition and maturation of ECM in the MTJs of murine forelimbs and (2) investigate musculoskeletal patterning and mechanical loading as contributing factors to MTJ morphogenesis. Combining 3D fluorescent imaging and quantitative proteomics, we demonstrated that ECM protein composition and specificity at the MTJ changed over development. Using mouse models in which muscle contractility (*mdg*) and patterning (*Tbx3* knockout) were disrupted, we found that static loading across the muscle-tendon interface appeared to be sufficient to initiate MTJ formation. Further maturation of the MTJ depended on cyclic loading; however, not insertion into an enthesis.

## Results

### The ECM undergoes distinct morphological and compositional changes during MTJ development

Due to the complex structure of the limb, methods for imaging cell and ECM distribution in 3D are needed to view structures that cannot be observed in traditional 2D cryosections [19]. Therefore, E12.5 – P21 murine forelimbs were visualized as either wholemount preparations, decellularized in sodium dodecyl sulfate (SDS) to remove light scattering lipids [20], or cleared using SeeDB [21] (**Figures 1A–1D**). Changes in MTJ morphology were assessed using antibodies against ECM enriched in muscle, MTJ, and tendon (**Figures 1E** and **1F)**.

**Figure 1:**
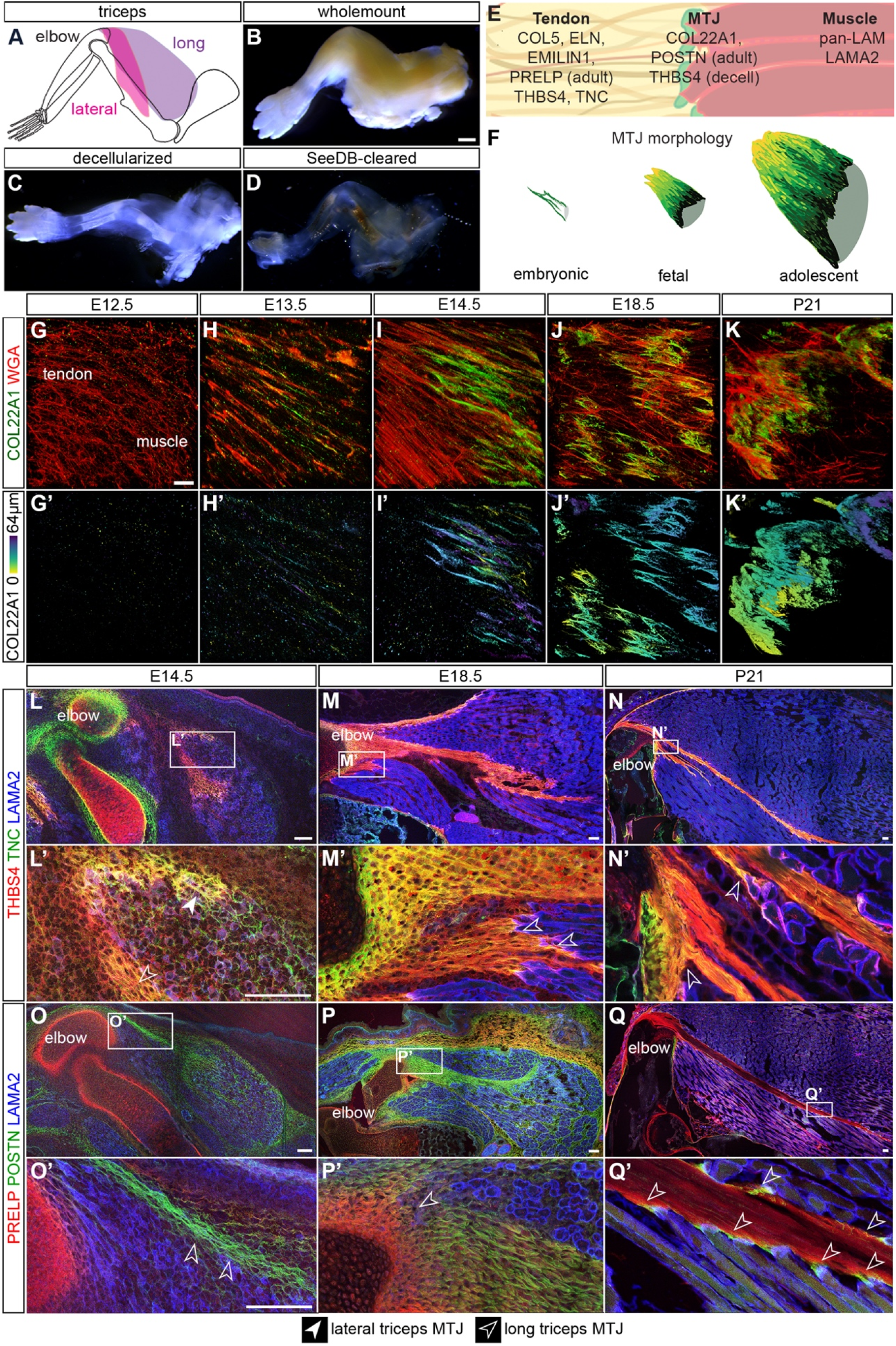
The ECM undergoes distinct morphological and compositional changes during MTJ development. **(A**) The insertion of the lateral triceps (pink) and long triceps (purple) into the olecranon (elbow) were either (**B**) wholemounted, (**C**) decellularized (decell) in SDS, or (**D**) SeeDB-cleared, stained with markers (**E**) used to differentially visualize muscle, MTJ, and tendon, then imaged. Representative E18.5 limbs shown. (**F**) Schematic of changes in MTJ morphology as assessed by COL22A1 staining. (**G-K’**) COL22A1 (green, depth projection) was identified at the putative MTJ starting at E13.5, and the morphology changed from thin, aligned structures at E13.5, to jagged protrusions at E18.5, then a highly interdigitated interface at P21. WGA (blue) marks general architecture. (**L-N’**) TNC (green) and THBS4 (red) were broadly distributed at E14.5 and became enriched in the tendon of the long (open arrowheads) and lateral triceps (closed arrowheads) at E18.5 and P21. (**O-Q’**) POSTN (green) was found in muscle (LAMA2, blue), tendon, and cartilage during development, but was restricted to the MTJ (arrowheads) at P21. PRELP (red) was found in the cartilage at E14.5 and 18.5, as well as the tendon at P21. G-K’: 63× decell, 3D rendering, z = 64 μm; L-Q: 10× cryosections; L’-Q’: 63× cryosections. Scale bars: 1 mm (B-D, L-Q); 10 μm (G-K’). Representative images from n = 3 biological replicates.

The presence of COL22A1, an ECM protein thought to physically link the basement membrane of individual myofibers with the tendon ECM [22], was negligible at E12.5 (**Figures 1G-G’**). By E13.5 - E14.5, COL22A1 was observed in linear structures between the developing lateral triceps tendon and muscle (**Figures 1H-1I’**). At E18.5, COL22A1 was organized into a cap-like structure with jagged protrusions (**Figures 1J** and **1J’**), and by P21 the cap contained shallow, serrated invaginations of greater frequency, but lower amplitude (**Figures 1K** and **1K’**). At all timepoints, COL22A1 was only found at the interface between muscle and tendon.

In contrast, other ECM varied in distribution across the muscle-tendon interface during development. Tenascin-C (TNC), a mechanically regulated matricellular protein [23], and THBS4, which is critical for collagen fiber spacing [24], were diffusely expressed in muscle and tendon at E14.5, then only found in the tendon at P21 (closed arrowhead = lateral triceps MTJ; open arrowhead = long triceps MTJ; **Figures 1L–1N’**). POSTN, a glycoprotein involved in collagen fiber regulation and crosslinking [25], was globally found in the muscle, MTJ, and tendon at E14.5 and E18.5; however, was restricted to the MTJ in the adult (**Figures 1O–1Q’**). PRELP, a small leucine-rich proteoglycan that binds both type I collagen (COL1) and basement membrane proteins [26], was only found in cartilage at E14.5 and E18.5, but the expression expanded to tendon in the adult (**Figures 1O–1Q’**). The differential dynamics of ECM distribution likely represents the changes needed to build and sustain a mechanically functional MTJ; however, the role of factors such as embryonic motility and muscle patterning on ECM deposition remains unclear.

### Muscle contraction was required for continuing development of the MTJ

To determine the role of embryonic motility in MTJ development, we first assessed when muscle contraction begins in the developing forelimb. Freshly explanted limbs were exposed to acetylcholine; muscle contraction in control limbs was observed at E13.5 and E14.5, but not at E12.5 (**Figures 2A–2C**; **Video S1**). We then tested the response of limbs from the *mdg* model, wherein a mutation in a Ca^2+^ channel subunit (CACNA1S) prevents muscle contraction [15, 16], and movement was absent as expected (**Figure 2D**; **Video S1**).

**Figure 2:**
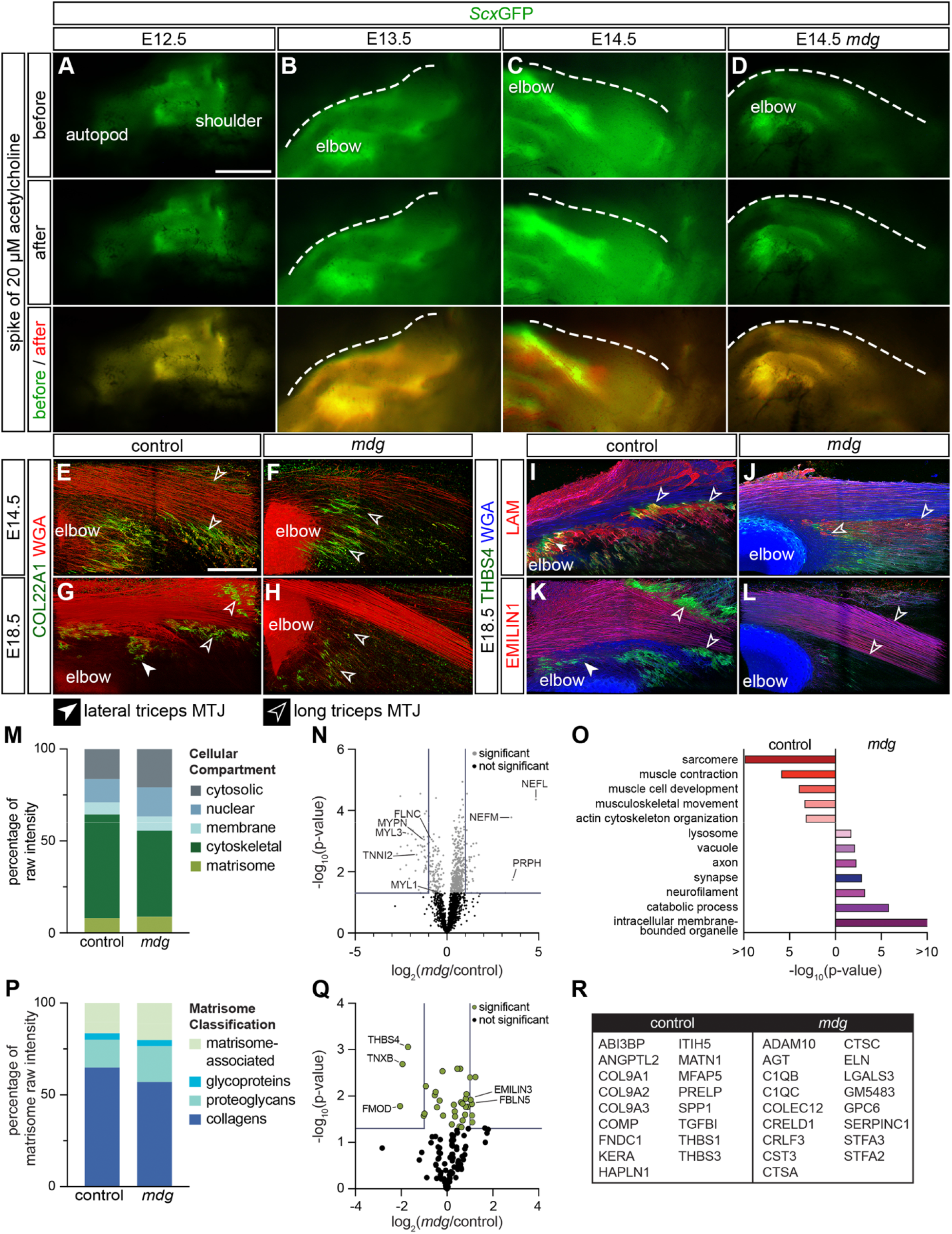
Muscle contraction was required for continuing development of the MTJ. (**A-D**) Distinct contraction of the triceps was observed in *Scx*GFP (green) forelimbs at E13.5 and E14.5 in response to acetylcholine, but not at E12.5 or in the *mdg* E14.5 limb. (**E-F**) The distribution of COL22A1 (green) was reduced at the termination of the long (open arrowheads) and lateral (closed arrowheads) triceps in *mdg* compared to control forelimbs at E14.5 (WGA, red). (**G-H**) By E18.5, COL22A1 was further reduced in the *mdg* mouse compared to control. (**I-J**) LAM^+^ muscle basement membranes (red) terminated in THBS4^+^ (green) MTJs in control, but not *mdg* limbs. WGA (blue) marks general architecture. (**K-L**) EMILIN1^+^ (red) long triceps tendons were reduced in size in the *mdg* limb. (**M**) Proteomic analysis of E18.5 triceps muscles revealed proteins from the cytoskeletal and nuclear compartment were significantly higher in *mdg* compared to controls (two-tailed t-test, cytoskeletal: p = 0.017; nuclear: p = 0.043; **Table S1**). (**N**) Volcano plot of the proteome in the *mdg* triceps versus control. Grey lines indicate ≥ 2-fold change and *p* < 0.05 (two-tailed t-test). (**O**) Selected gene ontology (GO) terms generated by analysis of proteins upregulated in, or exclusive to, the different genotypes. (**P**) The distribution of matrisome components was similar when muscle motility was disrupted (two-tailed t-test, all ns). (**Q**) Volcano plot of matrisome components. Grey lines indicate ≥ 2-fold change and *p* < 0.05 (two-tailed t-test). (**R**) Table of matrisome identified exclusively by LFQ in control and *mdg* triceps muscles. E-L: 63× decell, 3D rendering, z = 110 μm. Scale bars: 500 μm (A,D); 100 μm (E-L). A-L: representative images from n ≥ 3 biological replicates. M-R: average of n = 3 biological replicates.

We next assessed MTJ formation in *mdg* limbs and observed there was a slight reduction in COL22A1 at E14.5 where the tendon inserts into the long triceps muscle (**Figures 2E** and **2F**). By E18.5, THBS4 and COL22A1 were mostly absent in the MTJ of the *mdg* triceps (**Figures 2G** and **2H**). Additionally, there was a decrease in COL22A1 on the cartilage surface of E18.5 elbow and wrist joints within *mdg* limbs compared to controls (**Figures S1A-S1D’**). Tendon growth followed a similar pattern, where *mdg* and control tendons were of similar size at E14.5, but there was a marked decrease of EMILIN1 in *mdg* tendons at E18.5 (**Figures 2I–2L**). Comparable trends were observed at E14.5 and E18.5 in other *mdg* MTJs, including the extensor carpi ulnaris (ECU) in the wrist **(Figures S1E-S1M**).

To further investigate how muscle contraction contributes to the deposition of ECM proteins, or matrisome, we compared the proteome of triceps muscles harvested from E18.5 *mdg* and control mice using liquid chromatography-tandem mass spectrometry (LC-MS/MS). Overall, there was not a significant difference in the percentage of matrisome between genotypes; however, cytoskeletal proteins were significantly more abundant in controls compared to *mdg* muscles (**Figure 2M**). CACNA1S, the mutated protein in the *mdg* model, and striated muscle-related proteins, including CACNB1, CAPN3, MYL2, MYLK2, MYO18B, NRAP, and SMPX were exclusive to controls (**Table S1**), and FLNC, MYL1, MYL3, MYPN, and TNNI2 [27], were significantly reduced in *mdg* muscles (**Figure 2N**). Tenomodulin (TNMD), a tendon transcription factor [28], was also absent in the *mdg* triceps (**Table S1**). Gene ontology (GO) terms such as “sarcomere” and “muscle contraction” were generated by proteins enriched in, or exclusive to, control muscles. In contrast, terms related to muscle degradation, including “vacuole” and “catabolic processes” were generated by GO analysis for proteins enriched in, or exclusive to, *mdg* muscles (**Figures 2N** and **2O**; **Table S1**). Additionally, there was an elevation of axon-related proteins (NEFM, NEFL, PRPH) and neuron-related GO terms (“axon” and “neurofilament”) in *mdg* muscles (**Figures 2N** and **2O**), which corresponded to an expansion of neurofilament^+^ axons in the *mdg* limb (**Figures S1N-S1Q**).

There was a similar percentage of each matrisome classification in control and *mdg* triceps, and there was no difference in muscle-related basement membrane proteins such as LAMA2, NID1, and HSPG2 (**Figure 2Q**; **Table S1**). However, TNXB, FMOD, and THBS4 were significantly reduced in *mdg* limbs, and some ECM exclusive to controls (COMP, KERA, PRELP, THBS1, THBS3) were previously shown to be enriched in tendons [3] (**Figures 2Q** and **2R**). Together, the imaging and proteomic analyses indicated ECM content and organization were disrupted across the muscle-tendon interface of non-contractile *mdg* limbs.

### Knockout of *Tbx3* resulted in formation of an ectopic lateral triceps MTJ and tendon in E14.5 forelimbs

To investigate the role of muscle patterning in MTJ formation, we analyzed a model of ulnar mammary syndrome, a multi-system disorder caused by mutations in *TBX3* that results in abnormalities of the ulnar side of the limb [29]. Knockout of *Tbx3* in *Prrx1-expressing* connective tissue cells (*Prrx1Cre^Tg/+^;Tbx3^fl/fl^*) results in the lateral triceps ending in an ectopic insertion proximally within the long triceps muscle rather than at the olecranon [14].

At E14.5, the *Prrx1Cre^Tg/+^;Tbx3^fl/fl^* lateral triceps muscle was oriented perpendicular to the long axis of the humerus; however, the 3D organization of the fibrillar ECM (type V collagen, COL5) and vasculature (laminin, LAM) within the body of the muscle were maintained (**Figures 3A** and **3B**). When the matrisome of E14.5 forelimbs was analyzed using LC-MS/MS, the ECM composition was similar except for an elevation in TNC and TGFBI in *Prrx1Cre^Tg/+^;Tbx3^fl/fl^* compared with controls (**Figures 3C** and **3D**; **Table S2**). TNC and THBS4 were found in the ectopic insertion of the *Prrx1Cre^Tg/+^;Tbx3^fl/fl^* lateral triceps (arrows), as well as within the tendons of controls (**Figures 3E–3F’**). At E14.5, COL5, ELN, and EMILIN1 were also enriched at the insertions of both the long triceps tendon and the mispatterned lateral triceps (**Figures 3G–3L**). COL22A1^+^ fibers were observed at the ectopic insertion of *Prrx1Cre^Tg/+^;Tbx3^fl/fl^* limbs when rendered in 3D, but not the equivalent site of control limbs, suggesting that an ectopic MTJ formed (**Figures 3M–3N”**). The formation of the MTJ appeared delayed compared to controls, since COL22A1 was not observed in the ectopic triceps at E13.5 (**Figures S2A-S2D’**). *Scx*GFP^+^ cells were present in the lateral and long triceps tendons and colocalized with SOX9^+^ nuclei at the entheses in the elbow; however, *Scx*GFP was variably present at the ectopic insertion and did not colocalize with SOX9^+^ nuclei (**Figures 3O, 3P** and **S2E-S2I’**).

**Figure 3:**
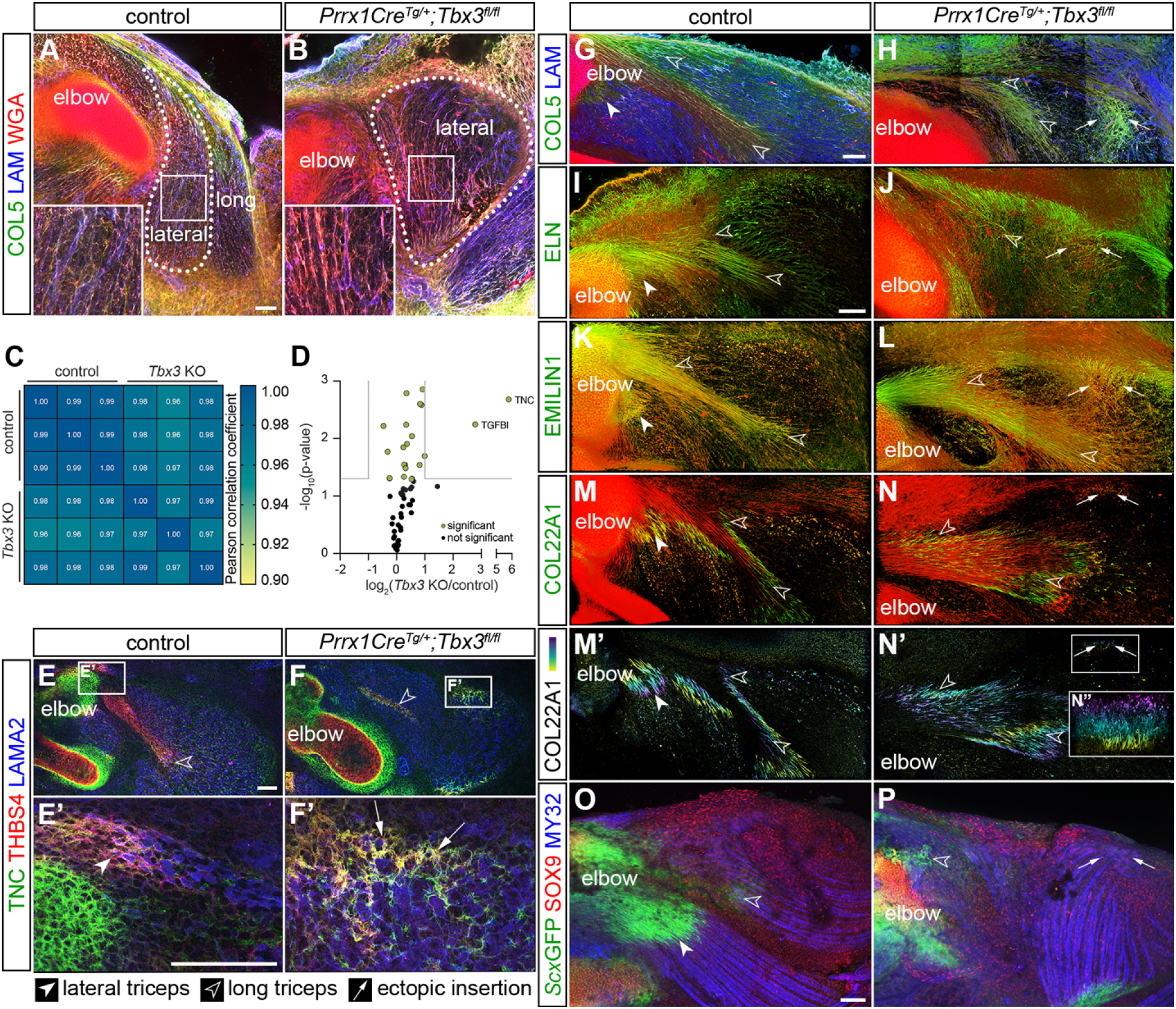
Knockout of *Tbx3* resulted in formation of an ectopic MTJ and tendon in E14.5 forelimbs. **(A, B**) Muscle fibers marked by COL5 (green) and WGA (red), and LAM^+^ (blue) blood vessels in the lateral triceps (dotted line) were morphologically normal, but abnormally oriented, in *Prrx1Cre^Tg/+^;Tbx3^fl/fl^* forelimbs. (**C**) Comparison of Pearson correlation coefficients revealed a high degree of similarity between the matrisome of *Prrx1Cre^Tg/+^;Tbx3^fl/fl^* forelimbs and controls. (**D**) Volcano plot of matrisome components. Grey lines indicate ≥ 2-fold change and *p* < 0.05 (two-tailed t-test); **Table S2**. (**E-F’**) The ectopic insertion of the LAMA2^+^ lateral triceps muscle (blue) was TNC^+^ (green) and THBS4^+^ (red). (**G-L**) COL5, ELN, EMILIN1 (green), and WGA (red) were enriched in tendons of the long and lateral triceps in the controls, as well as the ectopic insertion. (**M-N”**) COL22A1^+^ (green or depth projection) fibers marked the MTJ of the control lateral and long triceps insertions at the olecranon, as well as the ectopic insertion. Red = WGA. (**O, P**) SOX9^+^ (red) nuclei and *Scx*GFP^+^ (green) cells did not co-localize at the ectopic insertion of the MY32^+^ (blue) lateral triceps muscle, but were localized in control entheses. A, B: 10× decell, z projection, z = 19.2 μm, inset 2.5×; E, F: 10×, cryosections; E’, F’: 63× cryosections; G-N’: 25× decell, 3D rendering, z = 110 μm; N”: 3D rendering 90° rotation of ectopic insertion z = 175 μm; O, P: 10× wholemount, z projection, z = 458 μm. A-B, E-P: representative images from n = 3 biological replicates; scale bars: 100 μm. C-D: average of n = 3 biological replicates.

### The MTJ at the ectopic insertion was disorganized and lacked an enthesis at E18.5

The morphology of the E14.5 ectopic insertion suggested development of the MTJ, and potentially the enthesis, was delayed; therefore, we next looked at E18.5 forelimbs. In the control lateral head triceps muscle, LAM^+^ myofibers connected to a THSB4^+^ MTJ and converged into an EMILIN1^+^ tendon (**Figures 4A** and **4C**). At the ectopic insertion, LAM^+^ muscle transitioned into a disorganized region containing THBS4, EMILIN1, TNC, and *Scx*GFP (**Figures 4B–4D** and **4E-4H’’**). The distribution of these markers suggested the presence of an expanded, disorganized tendon, which was supported by a significant increase in TNC in the matrisome of the *Prrx1Cre^Tg/+^;Tbx3^fl/fl^* triceps (**Figure S3A**; **Table S3**).

**Figure 4:**
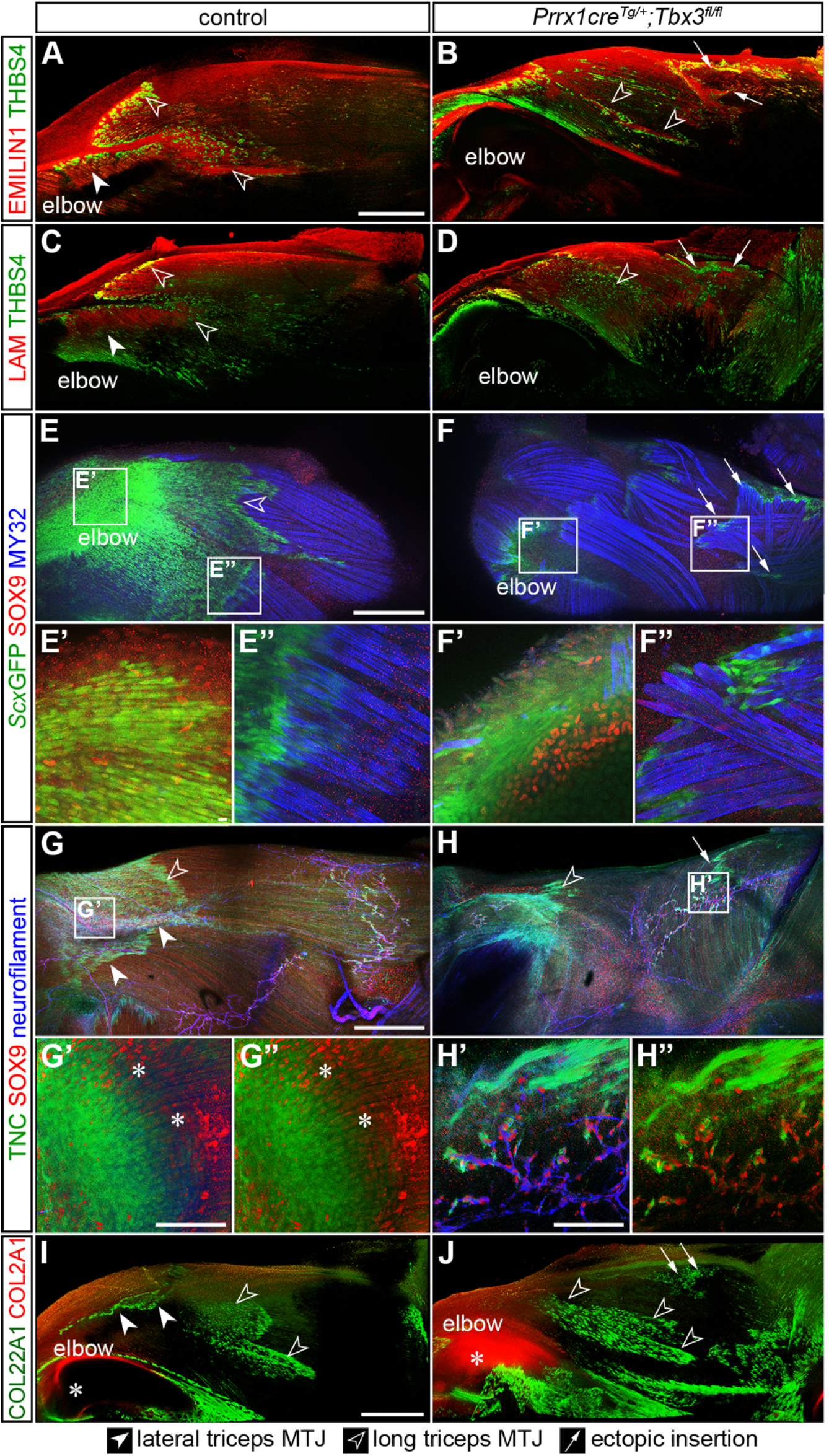
The MTJ at the ectopic insertion was disorganized and lacked an enthesis at E18.5. (**A-D**) THBS4^+^ (green) MTJs were present at the interface between LAM^+^ myofibers (red, A, B) and EMILIN1^+^ tendons (red, C, D) in the long and the lateral triceps in both control and *Prrx1Cre^Tg/+^;Tbx3^fl/fl^* limbs. (**E-F’’**) SOX9^+^ (red) nuclei co-localized with *Scx*GFP^+^ (green) cells at the enthesis of the control elbow (E’, F’); however, neither the control lateral triceps tendon nor the ectopic tendon contained SOX9^+^ nuclei (E’’, F’’). (**G-H’’**) SOX9^+^ nuclei (red) co-localized with TNC (green) and neurofilament^+^ (blue) neurons in both the control and *Prrx1Cre^Tg/+^;Tbx3^fl/fl^* triceps. SOX9^+^ nuclei and TNC co-localized (*) at the enthesis of the control elbow; however, the TNC^+^ ectopic tendon did not contain SOX9^+^ nuclei. (**I, J**) COL2A1 (red) was present in the cartilage of the humerus, but not in either the control or *Prrx1Cre^Tg/+^;Tbx3^fl/fl^* forelimbs in the area of the ectopic insertion marked by COL22A1 (green). A-D, I J: 10× decell, 3D rendering, z = 290 μm (A, B), z = 680 μm (C-D), z = 450 μm (I, J); E, F: 10× wholemount, z-projection, z = 40 μm; E-F’’: 63× wholemount, z-projection, z = 25 μm; G,H: 10× SeeDB-cleared, z-projection, z = 804 μm; G’-H’’ 63× 3D rendering, z = 86 μm. Scale bars: 500 μm (A-J); 10 μm (E, F); 100 μm (G, H). Representative images from n = 3 biological replicates.

Similar to E14.5, SOX9 and *Scx*GFP co-localized in cells at the insertion of the lateral and long triceps into the olecranon (**Figures 4E–4F’**). In contrast, SOX9^+^ nuclei did not colocalize with *Scx*GFP in the ectopic lateral triceps. However, SOX9^+^ nuclei were identified in arrays within the bodies of both control and ectopic lateral triceps muscles (**Figures 4E–4H’’**). These arrays were surrounded by TNC and contained neurofilament^+^ cells, suggesting that these SOX9^+^ nuclei were Schwann cells [30] (**Figures 4E–4H’’**). Notably, the nerves in *Prrx1Cre^Tg/+^;Tbx3^fl/fl^* forelimbs remained perpendicular to the long axis of the myofibers in the ectopic lateral triceps (**Figures S3B-S3G**). Furthermore, there was no deposition of COL2A1, an ECM component enriched in cartilage [31], in the proximity of the COL22A1^+^/TNC^+^ ectopic insertion in either decellularized or cryosectioned limbs (**Figures 4I, 4J,** and **S3H-S3I”**).

### The ectopic MTJ was mature by P21

To assess if mispatterning affected muscle and MTJ maturation, proteins from P21 triceps muscle-tendon units were analyzed using LC-MS/MS. There was a significant decrease in cytoskeletal components in *Prrx1Cre^Tg/+^;Tbx3^fl/fl^* triceps compared to controls (**Figure 5A**). Type I slow-twitch isoforms of muscle contractile proteins were significantly reduced (MYH7, MYL2, MYL3, TPM3, TNNI1, TNNT1, TNNC1) in *Prrx1Cre^Tg/+^;Tbx3^fl/fl^* triceps, but proteins associated with type II fibers (MYH2, MYH4) and embryonic/fetal myosins (MYH3, MYH8) were at the same relative abundance [27, 32] (**Figure 5B**; **Tables S4** and **S5**). GO analysis of proteins enriched in, or exclusive to, controls generated the terms “muscle contraction” and “transition between fast and slow fibers” (**Figure 5C**).

**Figure 5:**
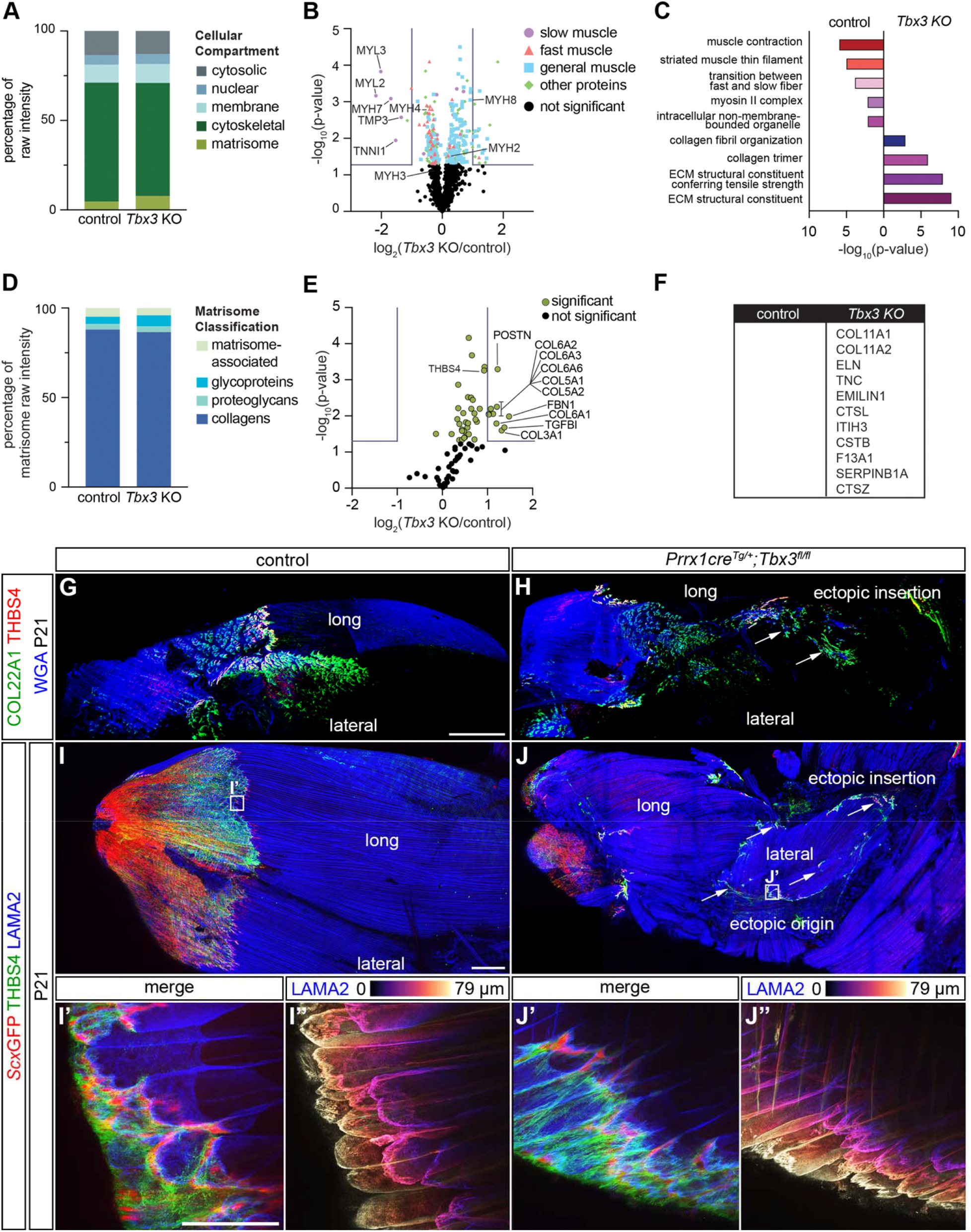
The ectopic MTJ was mature by P21. (**A**) Proteins from the cytoskeletal and matrisome fractions were significantly different in the P21 *Prrx1Cre^Tg/+^;Tbx3^fl/fl^* triceps unit compared with controls (two-tailed t-test: matrisome: p = 0.020; cytoskeletal: p < 0.0001; nuclear: p = 0.049; **Table S4**). (**B**) Volcano plot of all proteins revealed a reduction in slow-twitch muscle-related proteins. Grey lines indicate ≥ 2-fold change and *p* < 0.05 (two-tailed t-test). (**C**) Selected GO analysis terms for proteins upregulated, or exclusive to, the different genotypes. (**D**) The distribution of matrisome components was significantly different with more proteoglycan content in *Prrx1Cre^Tg/+^;Tbx3^fl/fl^* triceps (two-tailed t-test, proteoglycans: p = 0.027; all others: ns). (**E**) Volcano plot of matrisome components in control and *Prrx1Cre^Tg/+^;Tbx3^fl/fl^* triceps. Grey lines indicated ≥ 2-fold change and *p* < 0.05 (two-tailed t-test). (**F**) Matrisome proteins exclusive to control and *Prrx1Cre^Tg/+^;Tbx3^fl/fl^* forelimbs, as indicated by LFQ intensities. (**G-J**) COL22A1 (green) and THBS4 (red) were present in the MTJ of the control and ectopic lateral triceps. (**I’-J’’**) LAMA2^+^ (blue) muscle terminated in THBS4^+^ (green) and *Scx*GFP^+^ (red) tendons in control and ectopic lateral triceps MTJs. (**I’-J’**) LAMA2^+^ muscle fibers in both phenotypes terminated in morphologically mature MTJs. G, H: 10× decell, 3D rendering, z = 701 μm; I, J: 10× wholemount, z-projection, z = 1587 μm; I’-J’’: 63×, z-projection, z = 79 μm. Scale bars: 500 μm (G-J); 100 μm (I’ -J’’). A-F: average of n = 3 biological replicates. G-J”: representative images from n = 3 biological replicates.

There was a significant increase in the overall matrisome percentage and ECM proteoglycans in *Prrx1Cre^Tg/+^;Tbx3^fl/fl^* triceps (**Figures 5A** and **5D**). Also, tendon-related proteins including POSTN [33], THBS4 [24], EMILIN1 [34], TNC [35], COL11A1 [36], and COL11A2 [37], were exclusive to *Prrx1Cre^Tg/+^;Tbx3^fl/fl^* muscles (**Figures 5E–5F**). GO terms related to “ECM structural constituents” were only generated by proteins enriched in *Prrx1Cre^Tg/+^;Tbx3^fl/fl^* triceps (**Figure 5C**).

3D visualization of P21 forelimbs revealed both control and ectopic MTJs had a mature morphology and were COL22A1^+^ and THBS4^+^ (**Figures 5G–5J’’**). Furthermore, TNC and POSTN were enriched at the *Scx*GFP^+^ ectopic insertion at P21 (**Figure S4**). Together, this suggested the *Prrx1Cre^Tg/+^;Tbx3^fl/fl^* ectopic triceps muscle also included and MTJ and tendon.

### Muscle contraction was critical for the maturation of ectopic lateral triceps muscle and MTJ

Our results indicate that muscle contraction is necessary for MTJ maturation; however, the presence of COL22A1 and THBS4 at the *mdg* MTJ suggested the deposition of some ECM may also be influenced by static loading (**Figures 2E–2L**). The perpendicular orientation of the ectopic lateral triceps relative to the long axis of the humerus likely minimizes the contribution of static loading from longitudinal growth. Therefore, *Prrx1Cre^Tg/+^;Tbx3^fl/fl^;mdg* double knockouts were generated to remove both cyclic and static loads from the developing lateral triceps. As expected, muscle contraction was observed at E18.5 in both the control and *Prrx1Cre^Tg/+^; Tbx3^fl/fl^* ectopic lateral triceps, whereas the *mdg* and *Prrx1Cre^Tg/+^;Tbx3^fl/fl^;mdg* muscles did not contract (**Figures 6A–6D**; **Video S2**). The absence of contraction in *Prrx1Cre^Tg/+^;Tbx3^fl/fl^;mdg* limbs resulted in the loss of any discernable lateral triceps muscles, as well as the associated *Scx*GFP^+^/TNC^+^/THBS4^+^ tendons and COL22A1^+^ MTJs at both E16.5 and E18.5 (**Figures 6E–6L**, **S5**, and **S6**).

**Figure 6:**
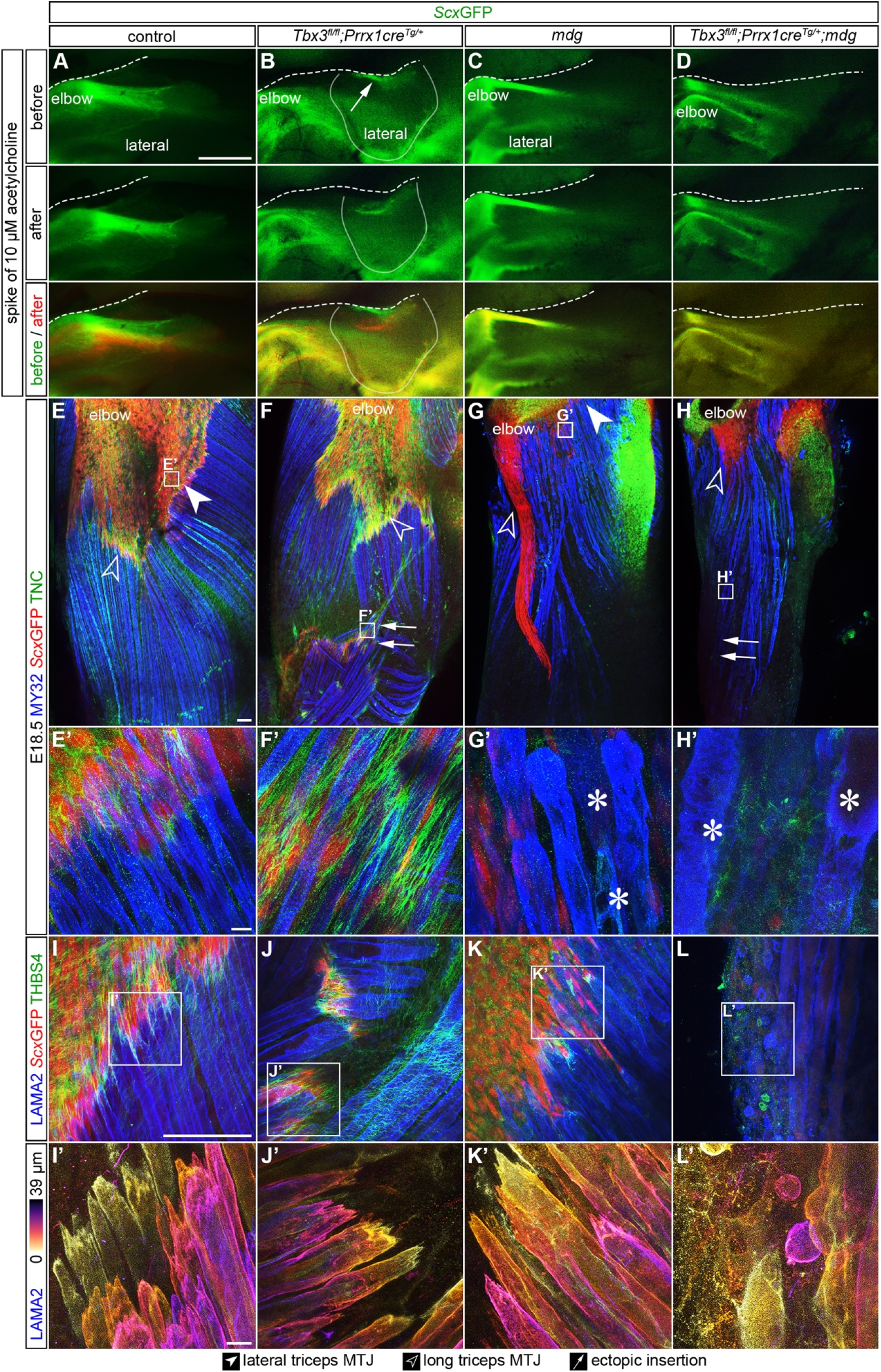
Muscle contraction was critical for the fetal maturation of ectopic lateral triceps muscle and MTJ. (**A-D**) Exposure of E18.5 limbs to 10 μm acetylcholine induced the contraction of the lateral triceps in control and *Prrx1Cre^Tg/+^;Tbx3^fl/fl^*, but not in *mdg* or *Prrx1Cre^Tg/+^;Tbx3^fl/fl^;mdg* limbs. Arrow indicates ectopic insertion. Dashed line indicates edge of tissue, solid line outlines the ectopic lateral triceps. (**E-H**) The lateral triceps was absent by E18.5 in the *Prrx1^CreTg/+^;Tbx3^fl/fl^;mdg* and TNC was not observed at the location of the putative ectopic insertion. (**E’-H’**) Striated muscle fibers were observed in the control and *Prrx1Cre^Tg/+^;Tbx3^fl/fl^* lateral triceps, but myofibers in the *mdg* and *Prrx1Cre^Tg/+^;Tbx3^fl/fl^;mdg* had reduced striations and contained vacuoles (*). (**I-L**) THBS4^+^ (green)/*Scx*GFP^+^ (red) tendons were present in the triceps of control, *Prrx1Cre^Tg/+^;Tbx3^fl/fl^*, and *mdg*, but not *Prrx1Cre^Tg/+^;Tbx3^fl/fl^;mdg* limbs. Blue = LAMA2 (muscle). (**I’-L’**) MTJs at the end of *mdg* myofibers were less mature, with fewer invaginations, than controls. E-H: 10× wholemount, z-projection, z = 547 μm; E’-H’, I-L’: 63× wholemount, z-projection, z = 44 μm (E’-H’) z = 39 μm (I, I’, K, K’, L, L’), z = 80 μm (J, J’). Scale bars: 500 μm (A-D); 100 μm (E-J); 10 μm (E’-J’). Representative images from n = 3 biological replicates.

Myofibers in control and *Prrx1Cre^Tg/+^;Tbx3^fl/fl^* lateral triceps were striated, but were disorganized and contained vacuoles (*) in *mdg* and *Prrx1Cre^Tg/+^;Tbx3^fl/fl^;mdg* muscles (**Figures 6E’–6H’**). Furthermore, the morphology of MTJs in *Prrx1Cre^Tg/+^;Tbx3^fl/fl^* and control limbs were similar, but the *mdg* and *Prrx1Cre^Tg/+^;Tbx3^fl/fl^;mdg* MTJs were less mature with fewer invaginations (**Figures 6I–6L’**). The triceps contained CD31^+^ blood vessels at a similar density as control limbs (**Figures S5**), suggesting vascular patterning was not disrupted in the absence of contractile muscle.

To assess if the lateral triceps formed, but regressed due to the lack of muscle contraction, an earlier embryonic timepoint was investigated. At E14.5, an ectopic lateral triceps was present in *Prrx1Cre^Tg/+^;Tbx3^fl/fl^;mdg* limbs; however, COL22A1, TNC, EMILIN1, and *Scx*GFP were absent at the location of the ectopic insertion (**Figures 7A–7H**, **S2G-S2I’**, and **S7**). Interestingly, the ECU and bicep brachii, which were not mispatterned in *Prrx1Cre^Tg/+^;Tbx3^fl/fl^;mdg* limbs and remained parallel to the direction of longitudinal growth, had a normal MTJ morphology (**Figure S7**). Additionally, the ECM fibers aligned in the direction of developing muscle fibers in control, *Prrx1Cre^Tg/+^;Tbx3^fl/fl^* and *mdg* limbs were no longer discernable in the *Prrx1Cre^Tg/+^;Tbx3^fl/fl^;mdg* ectopic lateral triceps (**Figure S7**; **Video S3**).

**Figure 7:**
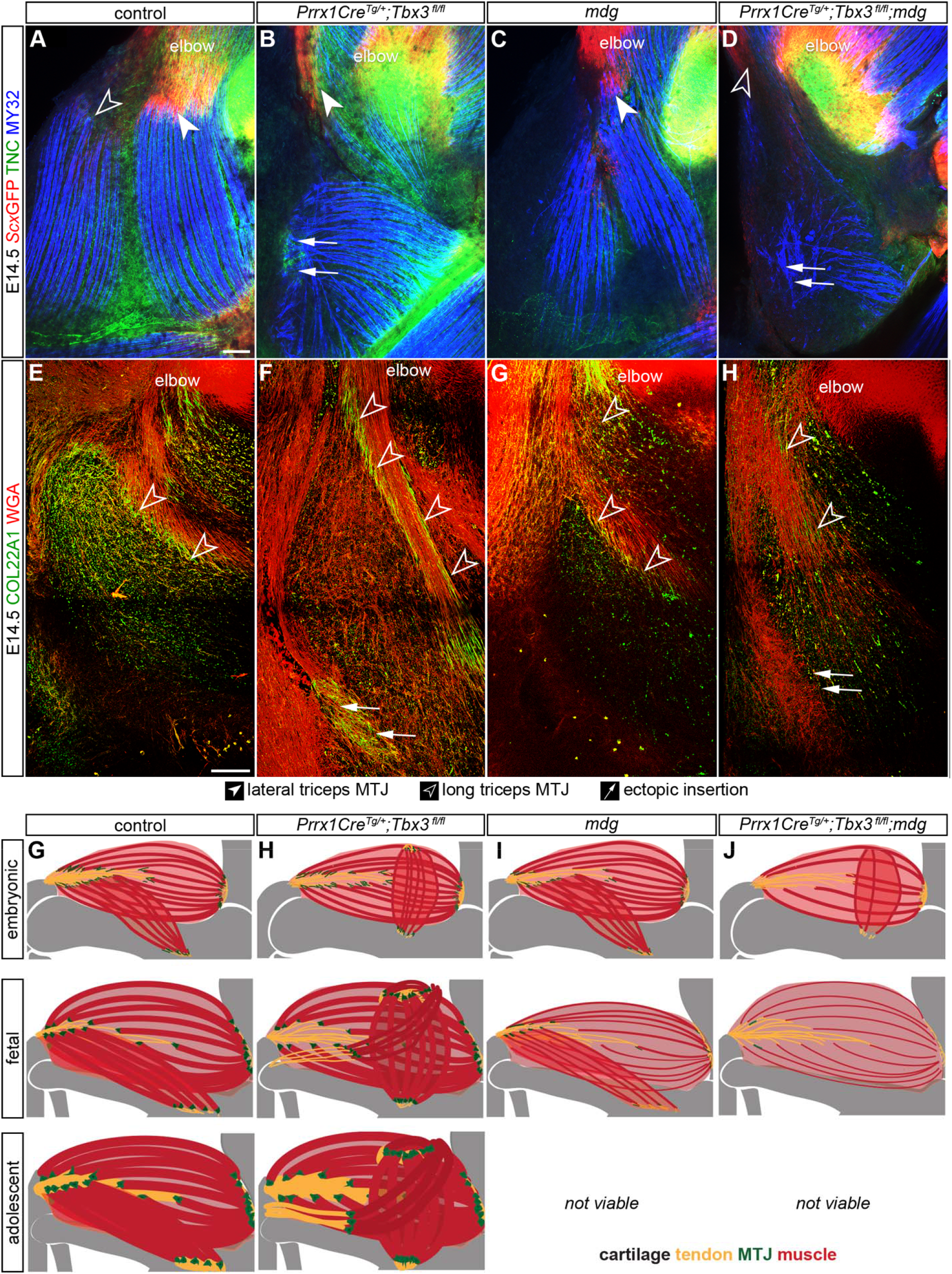
COL22A1 MTJ formation at the ectopic insertion depended on muscle contraction. (**A-D**) *Scx*GFP (green) and TNC (red) marked the long and lateral triceps tendon in E14.5 control, *mdg*, and *Tbx3* limbs. MY32 muscle (blue) did not terminate in *Scx*GFP^+^ or TNC^+^ tissue in the *Prrx1Cre^Tg/+^;Tbx3^fl/fl^;mdg* limbs. (**E-H**) At E14.5, COL22A1 (green) was present in the lateral and long triceps MTJ in both control and *Prrx1Cre^Tg/+^;Tbx3^fl/fl^*. COL22A1 was reduced in the *mdg* and absent in the ectopic insertion in *Prrx1Cre^Tg/+^;Tbx3^fl/fl^;mdg* limbs. (**G**) In the control lateral triceps, the organization of the COL22A1^+^ (green) MTJ developed from fibers (embryonic time point) to a cap-like structure with jagged protrusions (fetal), before maturing into caps with slight shallow serrated invaginations with a greater frequency, lower amplitude (adult). (**H**) When muscles were mispatterned (*Prrx1Cre^Tg/+^;Tbx3^fl/fl^*), the MTJ formed at the embryonic time points at the normal insertion of the long triceps and the ectopic insertion of the lateral triceps. (**I**) When muscle contraction was disrupted (*mdg*), the MTJ failed to mature. At embryonic time points, there was decreased COL22A1 in the MTJ, and by the fetal time point, the MTJ failed to mature, and tendon ECM decreased in abundance. (**J**) When mispatterned lateral triceps did not contract (*Prrx1Cre^Tg/+^;Tbx3^fl/fl^;mdg*), tendon and MTJ did not form in the embryonic limb, and by the fetal time point, the lateral triceps regressed. A-D: 10× wholemount, z-projection, z = 88 μm; E-H: 25× decell, 3D rendering, z = 78 μm. Scale bars: 100 μm. Representative images from n = 3 biological replicates.

## Discussion

Our results show the ECM in the developing MTJ undergoes dynamic changes in composition and morphology during development (**Figure 7G**). The maturation and specification of ECM at the MTJ depended on muscle contraction, as demonstrated using a murine model of muscle paralysis (*mdg*) (**Figure 7I**). We next investigated a model of musculoskeletal mispatterning (*Prrx1Cre^Tg/+^;Tbx3^fl/fl^*), where the lateral triceps was aligned perpendicular to the long axis of the humerus. This ectopic muscle contracted and inserted into a tendon at an MTJ, but there was no evidence of termination into an enthesis (**Figure 7H**). Removal of contraction in *Prrx1Cre^Tg/+^;Tbx3^fl/fl^;mdg* mice demonstrated the presence of the ectopic muscle and MTJ was regulated by muscle contraction, since both disappeared by E16.5 (**Figure 7J**). The misorientation of the lateral triceps also revealed that deposition of ECM at the MTJ is likely influenced by static loading as well since it was fully absent in the ectopic lateral triceps of *Prrx1Cre^Tg/+^;Tbx3^fl/fl^;mdg*, but not *mdg* limbs.

### Maturation of ECM at the MTJ depended on cyclic, and potentially static, loading

COL22A1 was first identified at the interface between muscle and tendon at E13.5-E14.5, organized in disperse, linear structures (**Figures 1G–1K’**). There was an increase in frequency and decrease in amplitude of the COL22A1^+^ cap-like structures between E13.5 and P21, consistent with the change in myofiber morphology observed with MTJ development [38]. These invaginations decrease stress at the interface by increasing the surface area, and are maintained at the MTJ by cyclic loading in the adult [39–42]. Forelimbs became responsive to acetylcholine between E12.5 and E13.5 (**Figures 2A–2D**), in line with previous reports that spontaneous limb twitching started between E13.0 and E14.0 [43], suggesting that cyclic loading is also important for MTJ development.

Since the deposition of COL22A1 coincided with the onset of contraction (**Figures 1G–1K’**), we analyzed E14.5 limbs in which muscles were paralyzed (*mdg*) to determine the direct influence of muscle contraction on the mammalian MTJ. We found there was no movement (**Figure 2D**; **Video S1**), even though distinct nerves already innervated the triceps unit (**Figures S3B-S3G**), and there was decreased COL22A1 in the *mdg* limb at the triceps MTJ (**Figures 2E–2F**). With extended loss of contractility, there was a dramatic reduction in COL22A1 within the long triceps MTJ at E16.5 and E18.5 and the joint surfaces (**Figures 2G–2H**, **S1A-S1D’**, **S5E-S5H**, and **S6I-S6L**), consistent with a prior study that reported the absence of MTJs in newborn *mdg* mice [44]. However, there was only a minor decrease in COL22A1 in MTJs of the *mdg* ECUs (**Figures S1E-S1F’**), which are located distal to the triceps, and may be more influenced by mechanical stimuli from movements by dam or wild type littermates [45].

The deposition of COL22A1 at the *Prrx1Cre^Tg/+^;Tbx3^fl/fl^* ectopic triceps insertion was delayed and was first observed at E14.5 (**Figures 3M–3N’** and **S2**), but obtained a mature morphology by P21, similar to the controls (**Figures 5I–5J’’**). The delay could be due to a decrease in static loading from the change in the angle of the ectopic lateral triceps insertion. Notably, COL22A1 was not found at the ectopic insertion of *Prrx1Cre^Tg/+^;Tbx3^fl/fl^;mdg* mice in which there was reduced static and cyclic loading (**Figures 7E–7H**). However, COL22A1 was observed in the MTJs of *Prrx1Cre^Tg/+^;Tbx3^fl/fl^;mdg* muscles that were still aligned in the direction of longitudinal growth in both the stylopod (biceps brachii) and on the ulnar side of the zeugopod (ECU) (**Figures S7A-S7H**). The influence of static loading has been observed in the development of other musculoskeletal tissues. Chick embryos with flaccid paralysis had a reduction in tendon equilibrium modulus, cross-sectional area, and ultimate tensile stress, while rigid paralysis did not have a significant effect on these properties [46], suggesting reduced static loading decreased tendon maturation.

COL22A1 is thought to play a role in mechanically linking the basement membrane that surrounds individual myofibers with the tendon ECM [22]. Recent evidence in zebrafish showed knockout of COL22A1 disrupted MTJ stability [47], suggesting presence of COL22A1 at the interface between muscle and tendon contributes to force transmission. The cellular origin of COL22A1 is currently unclear as it has been reported to be deposited by either muscle [5], myofibers after the fusion of fibroblasts during development [48], or a *Hic1^+^* mesenchymal progenitor [49]. Nevertheless, COL22A1 deposition depends on the presence of muscle, consistent with a previous study that investigated a mouse model in which limb muscles were absent (*Pax3^Cre/Cre^*) [20].

Likewise, expression of THBS4 in the tendon and MTJ was influenced by the presence and activity of muscle. Localization to the MTJ depended on muscle contraction (**Figures 2I–2L**), similar to zebrafish studies [7, 17], and THBS4 was no longer found in the tendons of muscle-less limbs [20]. THBS4 in decellularized tissues was only found at the MTJ (**Figures 2I–2L**), but was present throughout the tendon in cryosections (**Figures 1L–1N’**), suggesting it is more highly cross-linked or enriched at the MTJ [3, 50]. Given THBS4 regulates collagen fiber structure and ECM deposition [7, 24], the presence in the tendon and MTJ likely stabilizes the ECM at this interface to withstand muscle contraction.

### Maturation of muscle was affected by contraction and patterning

The ectopic lateral triceps in *Prrx1Cre^Tg/+^;Tbx3^fl/fl^* limbs appeared functional as the muscle contracted in response to acetylcholine and contained MY32^+^ striations indicative of maturing contractile machinery (**Figures 6E’–6H’**; **Video S2**). Interestingly, there was a specific reduction in type I slow-twitch fiber proteins in P21 *Prrx1Cre^Tg/+^;Tbx3^fl/fl^* muscles: MYH7, MYL2, MYL3, TPM3, TNNI1, TNNT1, and TNNC1 [27, 32] (**Figure 5B**; **Table 4**). This reduction could be due to defects in signaling from the knockout of *Tbx3* in connective tissue cells. For example, when *Tcf4^+^* connective tissue cells were knocked out, there were fewer slow-twitch fibers due to altered ECM or secreted factors [13]. Alternatively, the slow-to-fast muscle fiber type transition could result from reduced mechanical loading from the abnormal lateral triceps angle, which is known to influence fiber composition [51]

In contrast, myofibers in *mdg* muscles did not contract, had reduced striations and muscle cytoskeletal associated proteins, and contained vacuoles (**Figures 2N–2O** and **6E’–6H’**; **Table 1**), consistent with the autophagic vacuolar myopathy observed when CACNA1S is mutated [52]. When the lateral triceps was both mispatterned and non-contractile in *Prrx1Cre^Tg/+^;Tbx3^fl/fl^;mdg* limbs, there was a lack of any discernible lateral triceps myofibers (**Figures 6E–6H**, **S5A-S5D**, **S5I-S5L’**, and **S6A-S6H**). This may be due to regression of the muscle observed at E14.5 from the combined mispatterning (*Prrx1Cre^Tg/+^;Tbx3^fl/fl^*) and necrosis or autophagy of non-contractile *mdg* muscle fibers [53, 54].

### Tendon and MTJ in the ectopic insertion were disorganized but did not require insertion into an enthesis for maturation

To assess tendon formation at the ectopic insertion, we investigated the distribution of *Scx*, a transcription factor expressed in developing tendon and ligaments [55]. *Scx* is required for tendon elongation and growth [56], muscle attachment patterning [57], and enthesis development [58], and regulates the expression of COL1A1 and other later-stage tendon markers such as TNMD [59, 60]. We observed *Scx*GFP^+^ cells at the ectopic insertion as early as E14.5 and a distinct *Scx*GFP^+^ tendon at E16.5 and E18.5 (**Figures S2E-S2I’**, **S5A-S5D**, and **S6A-S6H**). The inconsistent presence of *Scx* at E14.5 was potentially due to variations in inter- and intra-litter maturation, as well as differences in *Prrx1Cre^Tg/+^;Tbx3^fl/fl^* phenotypes [14] (**Figures S2G-S2I’**). Alternatively, *Scx*GFP^+^ expression depends on mechanical loading, which was perturbed at the ectopic insertion as the muscle was no longer aligned with the axis of growth. Supporting this alternative, *Scx* and tendon-related gene expression increased in tendon cells after mechanical loading *in vitro* and increased after exercise *in vivo*, but decreased when muscle contraction was inhibited [61–64].

*Prrx1Cre^Tg/+^;Tbx3^fl/fl^* triceps contained elevated amounts of ECM associated with tendon such as EMILIN1, TNC, TGFBI, COL5A1, COL5A2, and COL11A1 [3]. However, staining for ECM enriched in the MTJ and tendon (COL22A1, THBS4, POSTN, and TNC) revealed the structures were disorganized (**Figures 3–5** and **S4-S7**). This disorganization could indicate the expression of ECM at this region was due to an injury response, rather than a reorientation of the tendon and MTJ. While POSTN and TNC are known to increase in fibrotic and injured muscle [65, 66], they were localized to regions also expressing *Scx*GFP at P21 (**Figures S4D-S4K’**). In contrast, only COL5 was observed throughout the interstitium of the ectopic muscle (**Figures S4J-S4K’**). Additionally, COL22A1 was not elevated in fibrotic tissue in late-stage muscular dystrophy models [67, 68]. Together, these data suggest that ectopic structures are tendon and MTJ, not a fibrotic response.

Although our data indicate the ectopic lateral triceps muscle terminates into a tendon with an MTJ, there was little evidence of an enthesis in this region as there was a lack of both cells that co-expressed SOX9^+^/*Scx*GFP^+^ and deposition of cartilage ECM (COL2A1) (**Figures 4E–4J** and **S3H-S3I’’**). Previous investigations of a skeletal-less model (*Prrx1Cre^Tg/+^;Sox9^fl/fl^*) suggested the presence of cartilage was necessary and sufficient to induce tendons in the autopod [10]. However, other studies indicated connective tissue patterning was decoupled from cartilage, suggesting a modularity in cartilage and tendon patterning for different limb regions (autopod, zeugopod, stylopod) [10, 69]. Data from the *Prrx1Cre^Tg/+^;Tbx3^fl/fl^* model suggests skeletal and connective tissue morphogenesis is decoupled in the stylopod and MTJ formation does not require associated skeletal patterning.

### Altered muscle patterning disrupts organization of ECM, nerve, and fatty tissue, but not blood vessels

Surprisingly, we observed minimal changes in early ECM composition of the forelimb and the patterning of the muscle interstitial ECM was normal at the scale of the myofibers. In the *Prrx1Cre^Tg/+^;Tbx3^fl/fl^* ectopic muscle, the interstitial ECM reoriented in the same direction as the myofibers (**>Figures 3A–3B**). However, the fibrillar ECM structures were not discernable in *Prrx1Cre^Tg/+^;Tbx3^fl/fl^;mdg* limbs, despite the presence of non-contractile myofibers (**Figures S7I-S7L**; **Video S3**). This suggested that when muscle mispatterning occurs, deposition of these ECM fibers requires reciprocal interactions between contractile muscle and connective tissue cells.

Similarly, the nerves reoriented along with the ectopic triceps in *Prrx1Cre^Tg/+^;Tbx3^fl/fl^* limbs. Other ectopic muscles were also reported to be innervated [70], supporting previous conclusions that nerve patterning involves factors intrinsic to the muscle [71, 72]. Notably, in limbs that did not contract, axons were expanded and mispatterned (**Figures S1N-S1Q’**) [73, 74]. In contrast, the blood vessels did not regress in *Prrx1^CreTg/+^;Tbx3^fl/fl^;mdg* limbs (**Figures S5I-S5L**). This is consistent with prior studies that suggested blood vessels form despite reduced metabolic requirements of the muscle [75]. Furthermore, in muscle-less limbs blood vessels form normally [20, 76].

Additionally, excess fatty/connective tissue was present at the surface of the ectopic lateral triceps in the *Prrx1Cre^Tg/+^;Tbx3^fl/fl^* limbs, which was also observed in an MRI of an ulnar-mammary syndrome patient [14]. The fatty accumulations could be due to an elevation in fibro/adipogenic progenitors, which are associated with mispatterning [12, 49].

Interestingly, the lateral triceps is also affected in other knockout models, including *Tbx5, Mox2*, and *Shox2* [11, 77, 78]. Knockout of *Shox2*, a proximal limb patterning transcription factor, disrupted lateral triceps structure resulting in a proximal insertion and abnormal gait that looked remarkably similar to the *Prrx1Cre^Tg/+^;Tbx3^fl/fl^* mice [14, 79]. This similarity is likely due to overlap in the affected areas: *Tbx3* alters the ulnar side while *Shox2* alters the proximal limb. In the *Shox2* knockout, nerve patterning was also disrupted likely due to mispatterned factors found in the disrupted lateral triceps, similar to the *Prrx1Cre^Tg/+^;Tbx3^fl/fl^* at E14.5 and older time points (**Figures S3B-S3I**) [79].

Overall, we observed the morphology of ECM at the MTJ matures from linear arrays at embryonic time points to a cap containing shallow, serrated invaginations of greater frequency, but lower amplitude in the postnatal forelimb. The maturation of the MTJ depended on muscle contraction, could occur when muscles were mispatterned, and was independent of enthesis formation.

## Experimental Procedures

### Transgenic mice and embryos

Wild type C57BL/6 mice were purchased from the Jackson Laboratory. *Tbx3^fl/fl^* [80] and *Prrx1Cre^Tg/+^* [81] mice were a generous gift from Drs. Anne Moon and Gabrielle Kardon. The *Scx*GFP [82] and *mdg* [15, 16] mice were generously provided by Dr. Ronen Schweitzer. Mice were maintained on a C57BL/6 background. Experimental protocols complied with, and were approved by, either the Purdue Animal Care and Use Committee (PACUC; protocol # 1209000723) or University of Colorado Boulder Institutional Animal Care and Use Committee (IACUC; protocol # 2705). PACUC and IACUC assess that Purdue University and University of Colorado Boulder researchers, respectively, and all procedures and facilities are compliant with regulations of the United States Department of Agriculture, United States Public Health Service, Animal Welfare Act, and university Animal Welfare Assurance. Mice were time mated, where embryonic day (E)0.5 refers to noon of the day when the copulation plug was noted. P21 mice and pregnant dams (E14.5, E18.5) were euthanized by CO_2_ inhalation and confirmed via cervical dislocation. E18.5 embryos were decapitated immediately after harvest. Embryos were transferred to chilled 1× PBS on ice for fine dissection of the forelimb. Tissues to be analyzed for LC-MS/MS were snap-frozen in liquid nitrogen and stored at −80°C until processed. Forelimbs for decellularization, SeeDB-clearing, and cryosectioning were processed immediately after harvest (as described below).

### Genotyping

Genotyping was performed using primers listed in **Table S1**. For proteomic experiments, *Prrx1Cre^tg/+^;Tbx3^fl/fl^* mice were used for *Tbx3* knockouts, and *Prrx1Cre^+/+^;Tbx3^fl/fl^* littermates were defined as controls. For imaging experiments, embryos with the *Prrx1Cre^+/+^* genotype were defined as controls. *mdg^+/+^* (control) and *mdg^+/−^* (heterozygotes) were used as controls and compared to *mdg^-/-^* littermates. For the muscle contraction studies and *Prrx1Cre^Tg/+^;Tbx3^fl/fl^;mdg* experiments, limbs were grouped via phenotype and validated by genotyping.

### Immunohistochemistry

For cryosectioning, forelimbs from embryos containing the *Scx*GFP transgene were fixed for 2 hours room temperature (RT) then overnight at 4°C. The samples were washed in 1× PBS, then incubated overnight at 4°C in sucrose solution (30% wt/vol solution of sucrose in PBS with 0.02% sodium azide). Subsequently, the samples were incubated in a 50% OCT:50% sucrose solution for 30 minutes. Samples were then embedded in optimal cutting temperature compound (OCT; Electron Microscopy Sciences), frozen in isopentane cooled dry ice, and stored at −80°C until cryosections were collected. *Scx*GFP negative samples were directly frozen. 10 μm cryosections were cut using a NX50 Cryostar Cryostat (Thermo Fisher Scientific), collected on charged slides and stored at −20°C until stained. Slides were stained via standard methods [3], using antibodies and probes at the concentrations indicated in **Table S1**. Negative controls consisted of the same processing, but with the exclusion of the primary antibodies. Samples were imaged on a Leica DMI6000 or Leica DM6 CFS STELLARIS upright confocal (Leica Microsystems). Images are representative of n = 3 biological replicates.

### 3D imaging of decellularized forelimbs

Forelimbs were processed for 3D imaging of the ECM using protocols modified from [20]. To improve tissue stability for early embryonic limbs, dissected forelimbs were embedded in 1% low melt agarose dissolved in Milli-Q water with 0.02% sodium azide. After solidifying at RT, agarose-embedded forelimbs were immersed in a solution containing sodium dodecyl sulfate (SDS). Samples were gently rocked at RT, with daily changes of the SDS solution for the times, concentrations, and volumes indicated in **Table S1**. After decellularization was complete, the samples were washed with 1× PBS for 1 hour at RT with rocking, fixed for 1 hour in 4% paraformaldehyde (PFA) in PBS at RT or overnight at 4°C with rocking, and washed with 1× PBS for 1 hour RT or 4°C overnight with rocking. The forelimbs were stored in 1× PBS at 4°C before processing for wholemount staining. The excess agarose was removed, and the isolated forelimbs were placed in a 48 or 96 well plate, blocked and permeabilized overnight at 4°C with 10% donkey serum and 0.02% sodium azide in 0.1% PBST (0.1% Triton X-100 in 1× PBS). The forelimbs were incubated with primary antibodies at concentrations indicated in **Table S1** in 0.2% bovine serum albumin (BSA) and 0.02% sodium azide in 1× PBS for 48 hours at 4°C with gentle rocking. After washing for 3 × 30 minutes in 0.1% PBST at RT, the forelimbs were incubated with fluorescently-labeled probes and secondary antibodies at concentrations listed in **Table S1**, diluted in 0.2% BSA and 0.02% sodium azide in 1× PBS at 4°C for 48 hours with gentle rocking while protected from light. The general structure of the limb was visualized by counterstaining with fluorophore-conjugated wheat germ agglutinin (WGA), which stains proteoglycans that contain sialic acid and n-acetylglucosamine [83]. After the forelimbs were washed 3 × 30 minutes in 0.1% PBST at RT, the samples were protected from light and stored in PBS at 4°C until imaged.

### Wholemount stained and SeeDB-cleared forelimbs

To compare the distribution of ECM with cells, limbs were wholemount stained and cleared using SeeDB following [21, 84]. In brief, forelimbs were washed in 1× PBS, fixed for 1 - 2 hours in 4% PFA in PBS at RT or overnight at 4°C with rocking. Forelimbs from embryos containing the *Scx*GFP transgene were fixed for 2 hours RT and overnight at 4°C. Samples were washed with 1× PBS for 1 hour RT or 4°C overnight with rocking and stored in 1× PBS at 4°C before processing for wholemount staining as described above (**Table S1**). For SeeDB-cleared tissues, samples were then incubated in a series of increasing concentrations of fructose diluted in 0.1× PBS and 0.5% thioglycerol. Specifically, forelimbs were incubated for 4-12 hours each in 20%, 40% and 60% fructose (weight/volume) for 12-24 hours each in 80% and 100%, and for 24-48 hours in 115% fructose. Sodium azide 0.1% was added to the 115% fructose solution for antifungal protection. Samples were stored in SeeDB 115% at RT until imaging.

### Imaging and image processing

Forelimbs were placed in an 8 well dish (ibidi) and either suspended in 1× PBS or embedded in a dish of low melt agarose and covered with 1× PBS. SeeDB-cleared limbs were placed in a custom PDMS imaging chamber with 115% fructose solution and covered with a coverslip. Forelimbs were imaged using either a Zeiss LSM 880 confocal microscope (Carl Zeiss Microscopy) with Zen 2.3 SP1 black software (V14.0.2.201) or Leica DM6 CFS STELLARIS upright confocal (Leica Microsystems) with LAS X software (V4.1.1.23273 to V4.4.0.24851). Confocal settings are indicated in **Table S1**. The laser power and gain were optimized for each sample to improve visualization of the ECM. Negative controls consisted of the same process without the addition of the primary antibody.

### Limb movement in response to acetylcholine

Limbs were isolated from freshly harvested embryos and the skin was gently removed prior to placing in a bed of 1% low melt agarose in a 100 mm dish. Once solidified, 50 mL of 1× PBS was added. 5 μL (E18.5) or 10 μL (E12.5-E14.5) of 100 mM stock acetylcholine was spiked in for a final concentration of 10 μM to 20 μM acetylcholine based on [85]. Movement was recorded by Infinity Analyze 7 via an Infinity 3 (Lumenera) camera on a Leica M80 microscope.

### Image processing

Confocal z-stacks of decellularized forelimbs were deconvoluted using either Zen Blue software (V2.3.64.0; Carl Zeiss Microscopy; nearest-neighbor deconvolution, settings: fast) or LAS X software (V4.1.1.23273 to V4.4.0.24851). To increase the signal intensity with depth if fluorescence intensity decayed, 10× and 25× decellularized tissues were bleach corrected (settings: exponential fit) [86]. Figures 3M–3N’ were processed with the despeckle filter, and visualized using FIJI 3D viewer [87]. Color coded stacks were rendered via the Temporal-Color Coder macro developed by Kota Miura at the Centre for Molecular and Cellular Imaging, EMBL Heidelberg, Germany. Cryosection images were processed with FIJI. Z-stacks of wholemount stained limbs were visualized via maximum z-projection. Renderings of forelimbs were compiled using Adobe Photoshop and Illustrator. Videos of muscle contraction were processed in Hitfilm Express.

### Proteomics analysis

E14.5 *Prrx1Cre^Tg/+^;Tbx3^fl/fl^* and control forelimbs were microdissected and processed as described in [3]. Briefly, forelimbs were homogenized, and intracellular components were removed by protein fractionation [3], leaving behind a matrisome-rich insoluble (IN) fraction. Previous work has shown that ECM proteins, especially from embryonic tissues, can be extracted into the CS fraction during the protein fractionation process [88]; thus, CS and IN fractions for E14.5 and E18.5 forelimbs were digested, cleaned [88] and analyzed by liquid chromatography tandem mass spectrometry (LC-MS/MS). Proteins were reduced, alkylated, and treated with 0.1U/200μL chondroitinase ABC for 2 hours before three enzymatic digestions: (1) 1μg LysC/200μL, 2 hours; (2) 3μg trypsin/200μL, overnight; and (3) 1.5μg trypsin/200μL, 2 hours. Samples were acidified to inactivate enzymes and peptides were cleaned with Pierce Detergent Removal Spin columns and C-18 MicroSpin columns. per the manufacturer’s protocol.

Triceps muscle-tendon units were isolated from E18.5 *Prrx1^CreTg/+^;Tbx3^fl/fl^* and control animals, and processed as described above. CS and IN fractions of mutant and control muscletendon units were analyzed by LC-MS/MS. Digested peptides from the E14.5 *Prrx1^CreTg/+^;Tbx3^fl/fl^* forelimbs and E18.5 *Prrx1^CreTg/+^;Tbx3^fl/fl^* muscle-tendon units were analyzed at the Purdue University Life Sciences Mass Spectrometry Facility. Samples were analyzed using the Dionex UltiMate 3000 RSLC Nano System coupled to the Q ExactiveTM HF Hybrid Quadrupole-Orbitrap Mass Spectrometer. Following digestion and clean up, 1μg of peptide was loaded onto a 300μm i.d. × 5mm C18 PepMapTM 100 trap column and washed for 5 minutes using 98% purified water/2% ACN/0.01% FA at a flow rate of 5 μL/minute. After washing, the trap column was switched in-line with a 75 μm × 50 cm reverse phase AcclaimTM C18 PepMapTM 100 analytical column heated to 50°C. Peptides were separated using a 120 minute gradient elution method at a flow rate of 300 nL/minute. Mobile phase A consisted of 0.01% FA in water while mobile phase B consisted of 0.01% FA in 80% ACN. The linear gradient started at 2% B and reached 10% B in 5 minutes, 30% B in 80 minutes, 45% B in 91 minutes, and 100% B in 93 minutes. The column was held at 100% B for the next 5 minutes before being brought back to 2% B and held for 20 minutes. Samples were injected into the QE HF through the Nanospray FlexTM Ion Source fitted with an emission tip. Data acquisition was performed monitoring the top 20 precursors at 120,000 resolution with an injection time of 100 ms.

Triceps units were isolated from E18.5 *mdg* and control, and P21 *Prrx1^CreTg/+^;Tbx3^fl/fl^* animals and tissues were processed as described in [3]. Whole muscles were homogenized in 8M urea, and proteins were digested and cleaned for LC-MS/MS as described above.

Peptides from the E18.5 *mdg* and P21 *Prrx1^CreTg/+^;Tbx3^fl/fl^* triceps were analyzed at the University of Colorado Boulder Proteomics and Mass Spectrometry facility using an UltiMate 3000 RSLC Nano System coupled to a QE HF-X. Before LC-MS/MS, peptides were cleaned up using a Waters Acquity M-class UV-UPLC with a rpC18 column and a fraction collector.

Raw files were processed by MaxQuant [89] (**Tables S1-S4)**. Data processing, statistical analysis, and visualization was done using Microsoft Excel and GraphPad Prism. Proteins were classified by either cellular compartment, matrisome classification [90, 91], or slow and fast muscle fiber type. A list of key muscle fiber protein types was compiled from the literature (**Table S4**). In addition, a list of proteins enriched in the different muscle fiber types were determined from [27]. For proteins with a statistically significant difference between fiber types, the fibers were annotated as slow if fiber type 1 was greater than 2a and 2x and fast if fiber type 1 was less than 2a and 2x. Proteins that did not meet this criteria or there was not a statistically significant difference were included as general. (**Table S4**) Raw intensities were used to calculate the distribution of Cellular Compartments and Matrisome Classifications. Log_2_-transformed LFQ intensities were used for volcano plot and Pearson correlation coefficient analyses.

GO analysis was performed on proteins significantly increased, or exclusive (via LFQ) to, P21 *Prrx1Cre ^Tg/+^;Tbx3^fl/fl^* vs control and E18.5 *mdg* vs control. Using g:profiler [92] and the list modified using REVIGO [93] using the setting: similarity: medium (0.7) and whole UniProt.

### Quantification and statistical analysis

Proteomics data were collected with n = 3 biological replicates and analyzed using Prism (GraphPad). The effect of genotype on cellular compartment and matrisome components were studied using two-tailed t-tests.

## Supporting information

Supplemental Images

## Abbreviations

MTJ, ECM

## Data availability statement

Mass spectrometry data supporting these results is openly available in the massIVE repository at MSV000089551. Reviewers please refer to cover letter for code.

## Acknowledgments

We would like to thank members of the Calve lab for assistance and helpful discussions. In particular, Drs. Yue Leng, Andrea Acuna, and Ye Bu for assistance with imaging and Dr. Karin Ejendal, Dr. Naagarajan Narayanan, Alexander Ocken, Caroline Hollier, Kristin Barringhaus, Madeline Ku, and Emmarie Ballard for help with mice and genotyping.

## Author contributions

S.N.L., K.R.J and S.C. designed the experiments; S.N.L., K.R.J., H.A.C, T.G.T., and K.P.M. performed the experiments; S.N.L., K.R.J., T.G.T., D.T.M., and S.C. analyzed the data; S.N.L. and S.C. interpreted the data and wrote the manuscript, with edits from K.R.J., H.A.C, T.G.T., D.T.M., and K.P.M.

