## Supplemental Images for "Mechanical loading is required for initiation of extracellular matrix deposition at the developing murine myotendinous junction"

**Video 1: Muscle contraction of the triceps in response to acetylcholine was observed as early as E13.5.** The skin was removed from *Scx*GFP$^{+}$ (green) E12.5 – E14.5 forelimbs and exposed to 20 µm acetylcholine. Movement was observed at E13.5 and later in control limbs but not in *mdg* limbs. Video is shown at 1× and 4× speed, forward (⏭) and reverse (⏮). Scale bars: 500 µm. Representative videos from n = 3 biological replicates.

**Video 2: The ectopic lateral triceps in *Prrx1$^{CreTg/+}$;Tbx3$^{fl/fl}$* limbs contracted in response to acetylcholine at E18.5.** The skin was removed from *Scx*GFP$^{+}$ (green) E18.5 forelimbs and exposed to 10 µm acetylcholine. Contraction was observed in the control and *Prrx1Cre$^{Tg/+}$;Tbx3$^{fl/fl}$* ectopic lateral triceps (arrows). No movement was detected in *mdg* or *Prrx1Cre$^{Tg/+}$;Tbx3$^{fl/fl}$;mdg* limbs. Video is shown at 1× and 4× speed, forward (⏭) and reverse (⏮). The end of the video shows control and *mdg* limbs in the same dish, exposed to the same spike of acetylcholine. Closed arrowheads: lateral triceps; open arrowheads: long triceps. Scale bars: 500 µm. Representative videos from n = 3 biological replicates.

**Video 3: EMILIN1+ fibers were absent from the belly of the ectopic lateral triceps muscle in *Prrx1Cre$^{Tg/+}$;Tbx3$^{fl/fl}$;mdg* limbs.** E14.5 decellularized forelimbs were stained for EMILIN1 (green), which marked fibers that were found within the lateral triceps of control, *mdg* (tendon: closed arrowhead) and *Prrx1Cre$^{Tg/+}$;Tbx3$^{fl/fl}$* (tendon: closed arrow) limbs. However, there were no fibers in the *Prrx1Cre$^{Tg/+}$;Tbx3$^{fl/fl}$;mdg* ectopic lateral triceps. Red: WGA; blue: THBS4. EMILIN1+ fibers were present in the long triceps (tendon: open arrowhead) of all genotypes. 10× decell, z = 114 µm. Scale bars: 100 µm. Representative videos from n = 3 biological replicates.

**Table S1**: Raw data supporting the E18.5 *mdg* versus control triceps muscle protein comparisons and methods details for primers, antibodies dilutions, decell protocol, and confocal settings.

**Table S2**: Raw data supporting the E14.5 *Prrx1Cre$^{Tg/+}$;Tbx3$^{fl/fl}$;mdg* versus control limb protein comparisons.

**Table S3**: Raw data supporting the E18.5 *Prrx1Cre$^{Tg/+}$;Tbx3$^{fl/fl}$;mdg* versus control triceps protein comparisons.

**Table S4:** Raw data supporting the P21 *Prrx1Cre$^{Tg/+}$;Tbx3$^{fl/fl}$;mdg* versus control triceps protein comparisons and slow verse fast muscle classification support.

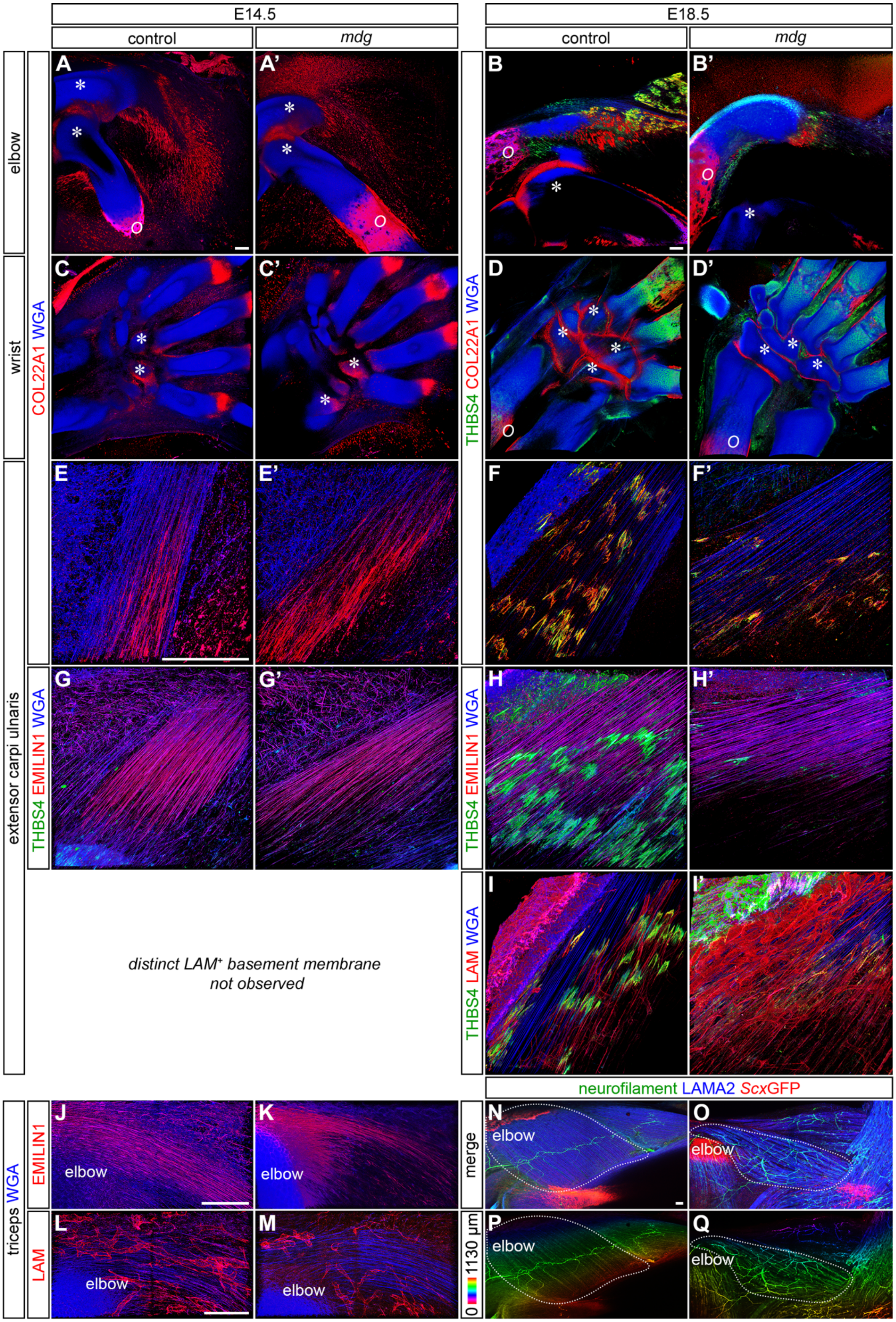

E14.5
E18.5
control
mdg
control
mdg
elbow
wrist
extensor carpi ulnaris
COL22A1 WGA
THBS4 COL22A1 WGA
THBS4 EMILIN1 WGA
THBS4 LAM WGA
distinct LAM+ basement membrane
not observed
triceps WGA
EMILIN1
LAM
merge
0 1130 µm
neurofilament LAMA2 ScxGFP
elbow

**Figure S1*: *Muscle contraction was required for MTJ maturation in the wrist, expansion of COL22A1 at the joint surface, and nerve patterning.** (**A, A', C, C**') COL22A1 (red) was expressed on joint surfaces (*) within the elbow and wrist in both control and *mdg* limbs at E14.5 (WGA, blue). (**B, B', D, D'**) COL22A1 was expanded around the humerus and ulna (B) and wrist bones (D) joint surfaces (*) in E18.5 controls, but not in the *mdg* limbs. Ossification centers (*O*) were still present. (**E, E', G-G'**) COL22A1 and EMILIN1 (red) were observed in the E14.5 extensor carpi ulnaris (ECU) in both control and *mdg* limbs. THBS4 (green) was not MTJ-specific at E14.5. (**F, F', H, H', I, I'**) COL22A1 (red) and THBS4 (green) were reduced at the MTJ of *mdg* E18.5 ECU when compared to controls. (**J, K**) EMILIN1$^+$ tendons (red) were slightly reduced in thickness in the *mdg* triceps. (**L, M**) LAM$^+$ (red) blood vessel patterning was similar in control and *mdg* triceps. (**N-Q**) The distribution of neurofilament$^+$ nerves (green or depth projection) was expanded in LAMA2$^+$ (blue) *mdg* triceps muscles (dotted line) compared to controls. Tendons labeled by *Scx*GFP (red). A-D': 10× decell, 3D rendered, z = 342 µm (A-B'), z = 496.2 µm (C, C', D, D'); E-F', H-M: 63× decell, 3D rendered, z = 65 µm (E-F', H-I'), z = 59 µm (G,G'), z = 102 µm (J, K), 80 µm (L,M); N-Q: 10× wholemount, z-projection, z =1130 µm. Scale bars: 100 µm. Representative images from n = 3 biological replicates.

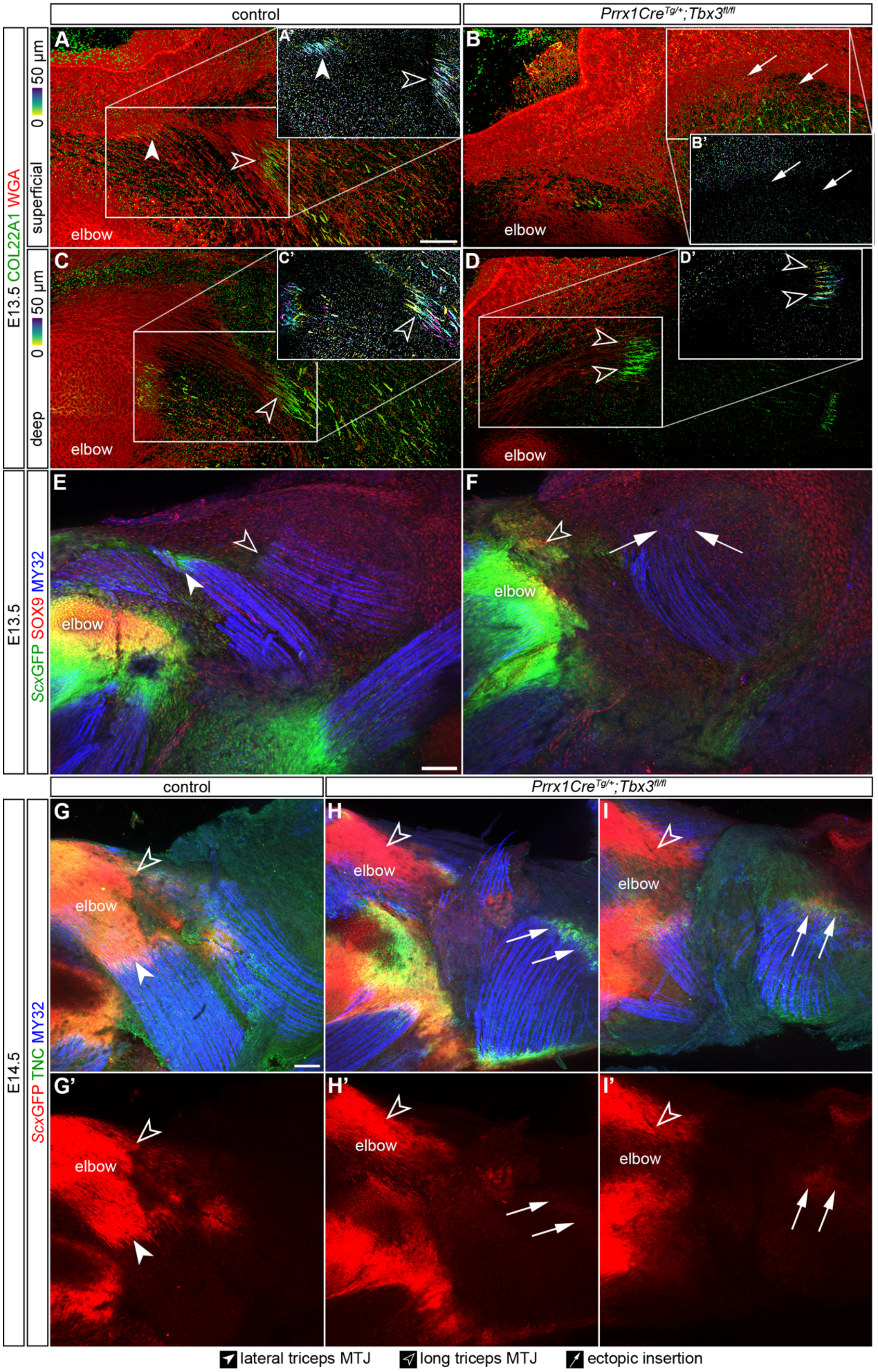

control
Prrx1Cre^Tg/+;Tbx3^fl/fl
E13.5 COL22A1 WGA
superficial
deep
0 50 µm
elbow
E13.5
ScxGFP SOX9 MY32
E14.5
ScxGFP TNC MY32
lateral triceps MTJ
long triceps MTJ
ectopic insertion

**Figure S2: Development of the MTJ, tendon, and cartilage at E13.5-E14.5 in *Prrx1Cre*$^{Tg/+}$*;Tbx3*$^{fl/fl}$ limbs.** (**A-D'**) At E13.5, COL22A1 (green or depth projection) was deposited at the MTJ of the lateral and long triceps. No discernable COL22A1 was observed at the ectopic insertion. **(E-F)** *Scx*GFP (red) and SOX9 (green) did not co-localize at the ectopic insertion at E13.5, but did co-localize at the enthesis of the normal insertion. MY32 = muscle (blue). (**G-I'**) *Scx*GFP$^{+}$ (red) cells were inconsistently observed at the TNC$^{+}$ (green) ectopic insertion at E14.5, and was not observed in the same area within control limbs. MY32 = muscle (blue). A-D': 25× decell, 3D rendering, z = 50 µm (A-B), z = 56 µm (C-D); E, F: 10× wholemount, z-projection, z = 458 µm (E, F); G-H': 10× wholemount, z-projection, z = 381.6 µm. Scale bars: 100 µm. Representative images from n = 3 biological replicates.

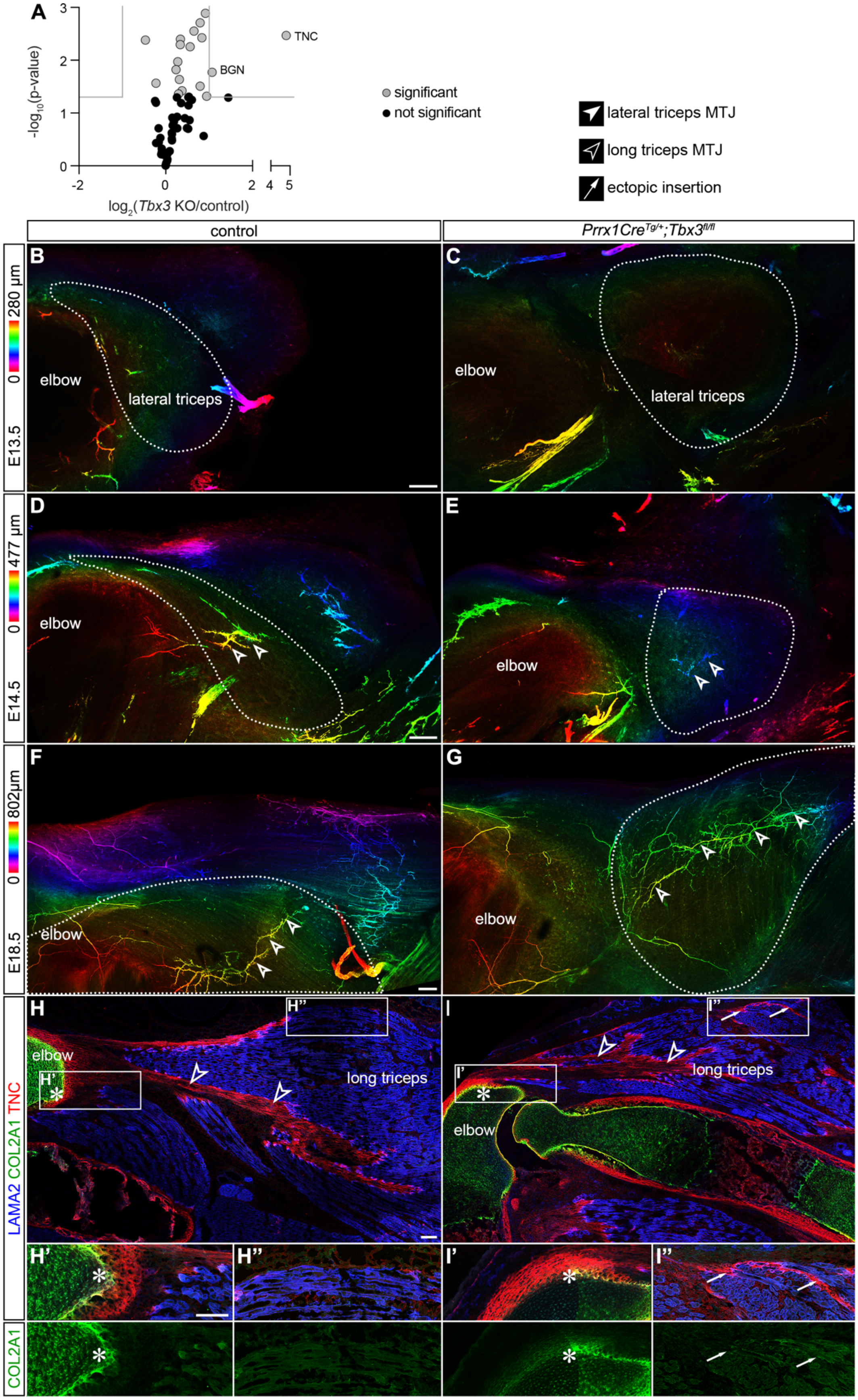
A
TNC
BGN
-log10(p-value)
log2(Tbx3 KO/control)
significant
not significant
lateral triceps MTJ
long triceps MTJ
ectopic insertion
control
Prrx1Cre^Tg/+;Tbx3^fl/fl
B
C
D
E
F
G
H
I
H'
H''
I'
I''
E13.5
0
280 µm
E14.5
477 µm
E18.5
802µm
elbow
lateral triceps
long triceps
LAMA2 COL2A1 TNC
COL2A1

**Figure S3: Development of the nerve and cartilage from E13.5 – E18.5 in *Prrx1Cre$^{Tg/+}$;Tbx3$^{fl/fl}$* limbs. (A)** Volcano plot of matrisome components identified in control and *Prrx1Cre$^{Tg/+}$;Tbx3$^{fl/fl}$* E18.5 triceps forelimbs (**Table S3**). Grey lines indicate ≥ 2-fold change and $p < 0.05$ (two-tailed t-test). Average of n = 3 biological replicates. (**B-G**) Neurofilament$^{+}$ (depth projection) nerves (arrowheads) were not observed at E13.5 in either the *Prrx1Cre$^{Tg/+}$;Tbx3$^{fl/fl}$* or control muscles. At E14.5 and E18.5, neurofilament$^{+}$ nerves were observed within the lateral triceps (dotted line), and were oriented perpendicular to the direction of muscle fibers, even in the mispatterned *Prrx1Cre$^{Tg/+}$;Tbx3$^{fl/fl}$* limbs. (**H-I''**) While overlap between the TNC$^{+}$ (red) tendon and COL2A1$^{+}$ (green) cartilage was observed in the normal enthesis at E18.5 (*), COL2A1 did not co-localize with the TNC$^{+}$ ectopic insertion (arrows). LAMA2 = muscle (blue). B-G: 10× wholemount (B, C), SeeDB-cleared (D-G), z-projection, z = 280 µm (B, C), z = 427 µm (D, E), z = 802 µm (F, G); H-I'': 10× cryosections (H, I); 63×, cryosections (H', H'', I', I''). Scale bars: 100 µm. Representative images from n = 3 biological replicates.

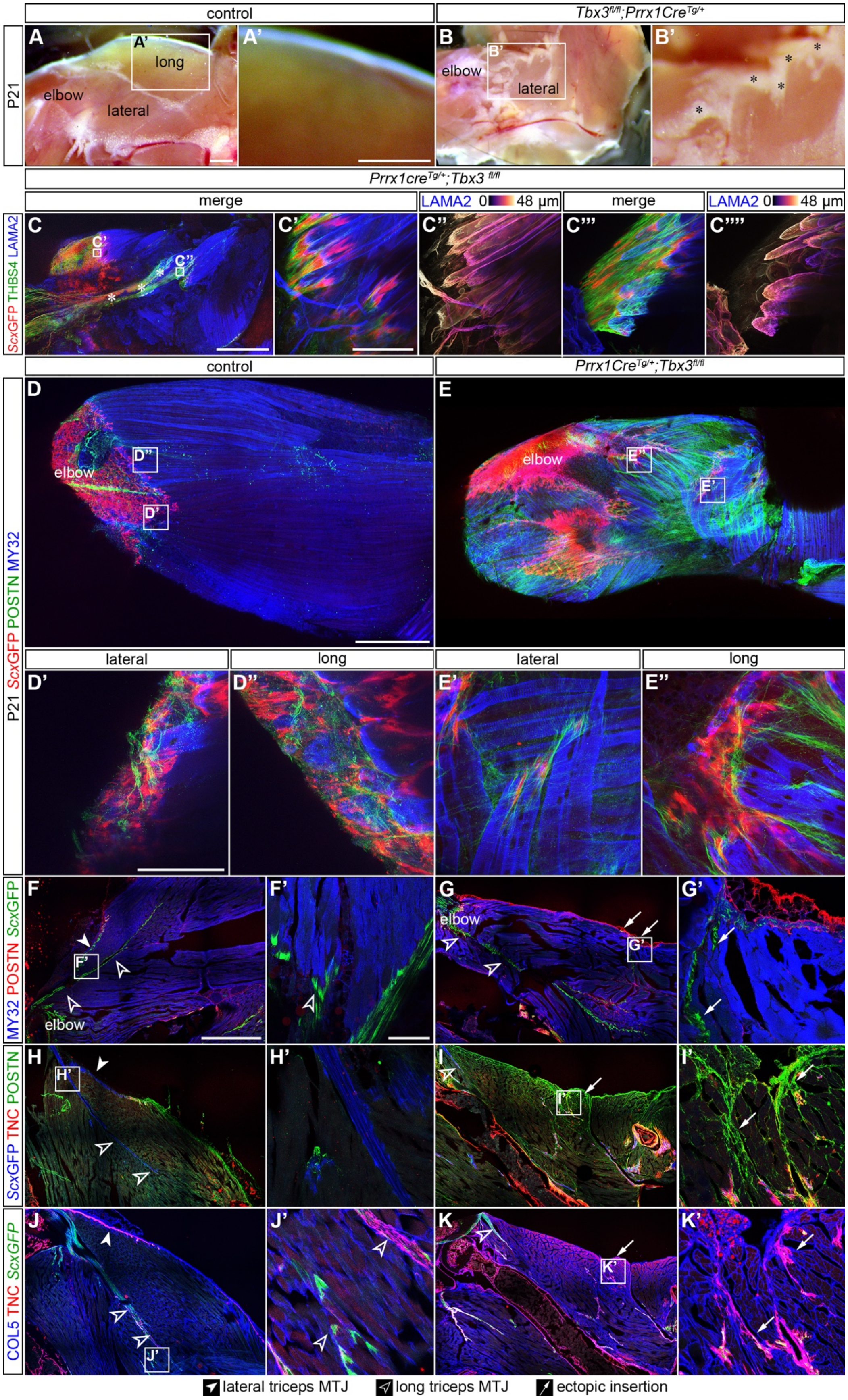
control
Tbx3fl/fl;Prrx1CreTg/+
P21
A
A'
long
elbow
lateral
B
B'
Prrx1creTg/+;Tbx3 fl/fl
merge
LAMA2 0 48 µm
ScxGFP THBS4 LAMA2
C
C'
C''
C'''
C''''
Prrx1CreTg/+;Tbx3fl/fl
D
D'
D''
E
E'
E''
P21 ScxGFP POSTN MY32
MY32 POSTN ScxGFP
F
F'
G
G'
ScxGFP TNC POSTN
H
H'
I
I'
COL5 TNC ScxGFP
J
J'
K
K'
lateral triceps MTJ
long triceps MTJ
ectopic insertion

**Figure S4: There was increased fat, POSTN, and TNC accumulation in the *Prrx1$^{CreTg/+}$; Tbx3$^{fl/fl}$* ectopic triceps.** (**A-B'**) The *Prrx1Cre$^{Tg/+}$;Tbx3$^{fl/fl}$* lateral triceps had increased fat (*) compared to the controls. (**C-C''''**) In some P21 limbs, it appeared an ectopic origin was also present, connected to the elbow by a THBS4$^{+}$/*Scx*GFP$^{+}$ tendon (*). Both the long triceps insertion and ectopic triceps origin muscle fibers terminated in a mature MTJ. (**D-K'**) POSTN and TNC were enriched near the *Scx*GFP+ ectopic insertion. COL5 was found thorough the interstitial spaces surrounding the muscle. C: 10× wholemount, z-projection, z = 1687 µm I; z = 1160 µm (D, E), C'-C'''' 63× wholemount, z-projection, z = 48 µm (C'-C'''), z = 51 µm (D', D'', E', E''), F-K: 10× cryosections, F'-K': 10× cryosections. Scale bars: 1 mm (A-B', C-K); 100 µm (C'-C''', D', D'', E', E'', F'-K'). Representative images from n = 3 biological replicates.

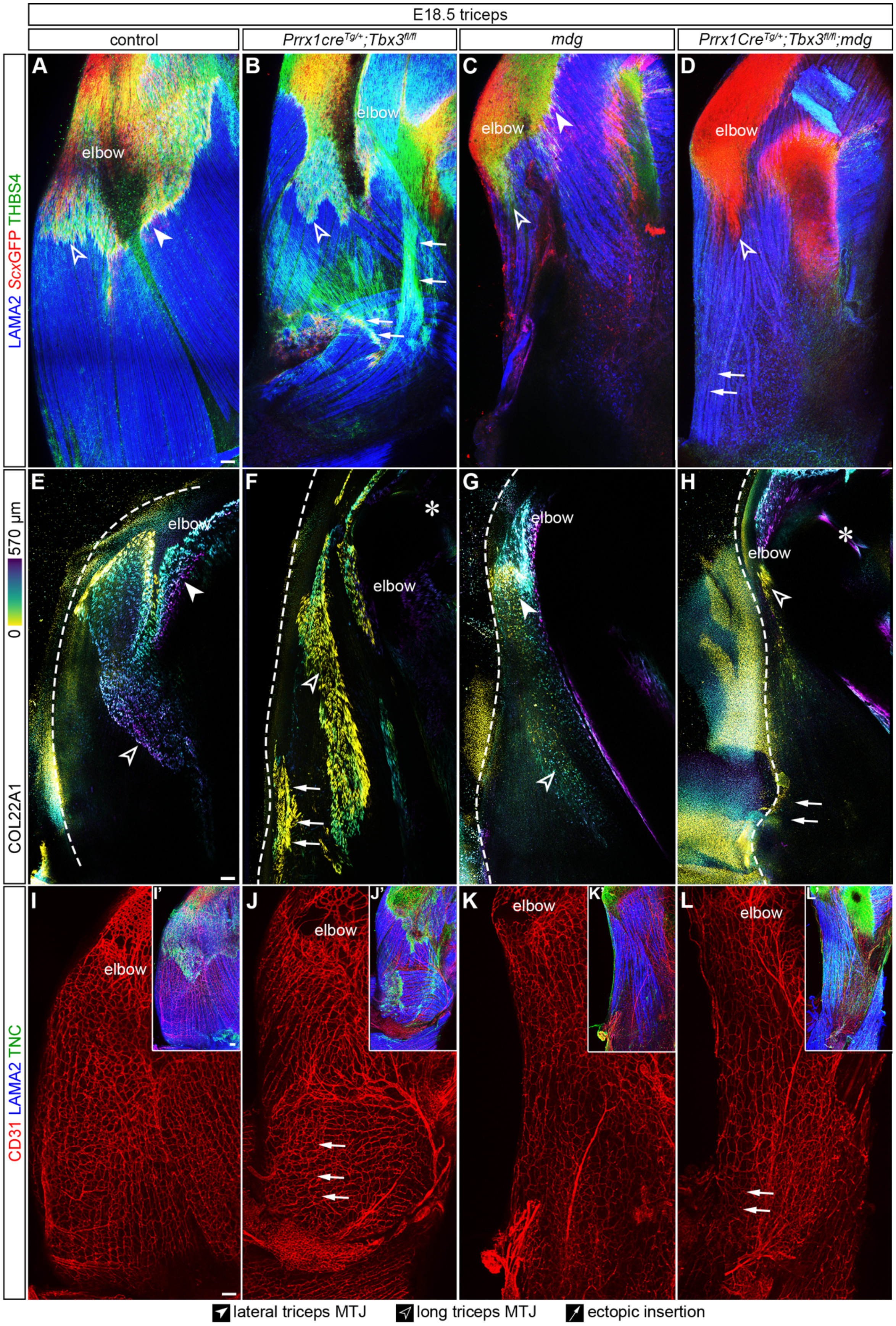

E18.5 triceps
control
Prrx1cre^Tg/+;Tbx3^fl/fl
mdg
Prrx1Cre^Tg/+;Tbx3^fl/fl;mdg
LAMA2 ScxGFP THBS4
COL22A1
0
570 µm
CD31 LAMA2 TNC
A
B
C
D
E
F
G
H
I
I'
J
J'
K
K'
L
L'
elbow
lateral triceps MTJ
long triceps MTJ
ectopic insertion

**Figure S5: The ectopic lateral triceps muscle, MTJ, and tendon were not observed in E18.5 *Prrx1Cre$^{Tg/+}$;Tbx3$^{fl/fl}$;mdg* limbs, but blood vessel structure was maintained.** (**A-D**) LAMA2$^{+}$ (blue) lateral triceps muscle and THBS4$^{+}$ (green)/*Scx*GFP$^{+}$ (green) tendon formed in the proper location in control and *mdg* limbs. The ectopic lateral triceps muscle was present in *Prrx1Cre$^{Tg/+}$;Tbx3$^{fl/fl}$* but not *Prrx1Cre$^{Tg/+}$;Tbx3$^{fl/fl}$;mdg* limbs. (**E-H**) COL22A1$^{+}$ (depth projection) was present in the lateral and long triceps and ectopic insertion. In contrast, COL22A1 was weakly found at the MTJ of the long triceps tendon and not found at the location of the ectopic insertion in *Prrx1Cre$^{Tg/+}$;Tbx3$^{fl/fl}$;mdg* limbs. * Indicates joint surface. (**I-L'**) CD31$^{+}$ (red) blood vessels were present in the location of the lateral triceps in *mdg* and *Prrx1Cre$^{Tg/+}$;Tbx3$^{fl/fl}$;mdg* limbs. TNC = green; MY32 = blue. A-D, I-L': 10× wholemount, z-projection, z = 387 µm (A, C, D), z = 501 µm (B), z = 497 µm (I), z = 1130 µm (J), z = 1009 µm (K), z = 602 µm (L); E-H: 10× decell, 3D rendering, z = 570 µm (E-H). Scale bars: 100 µm. Representative images from n = 3 biological replicates.

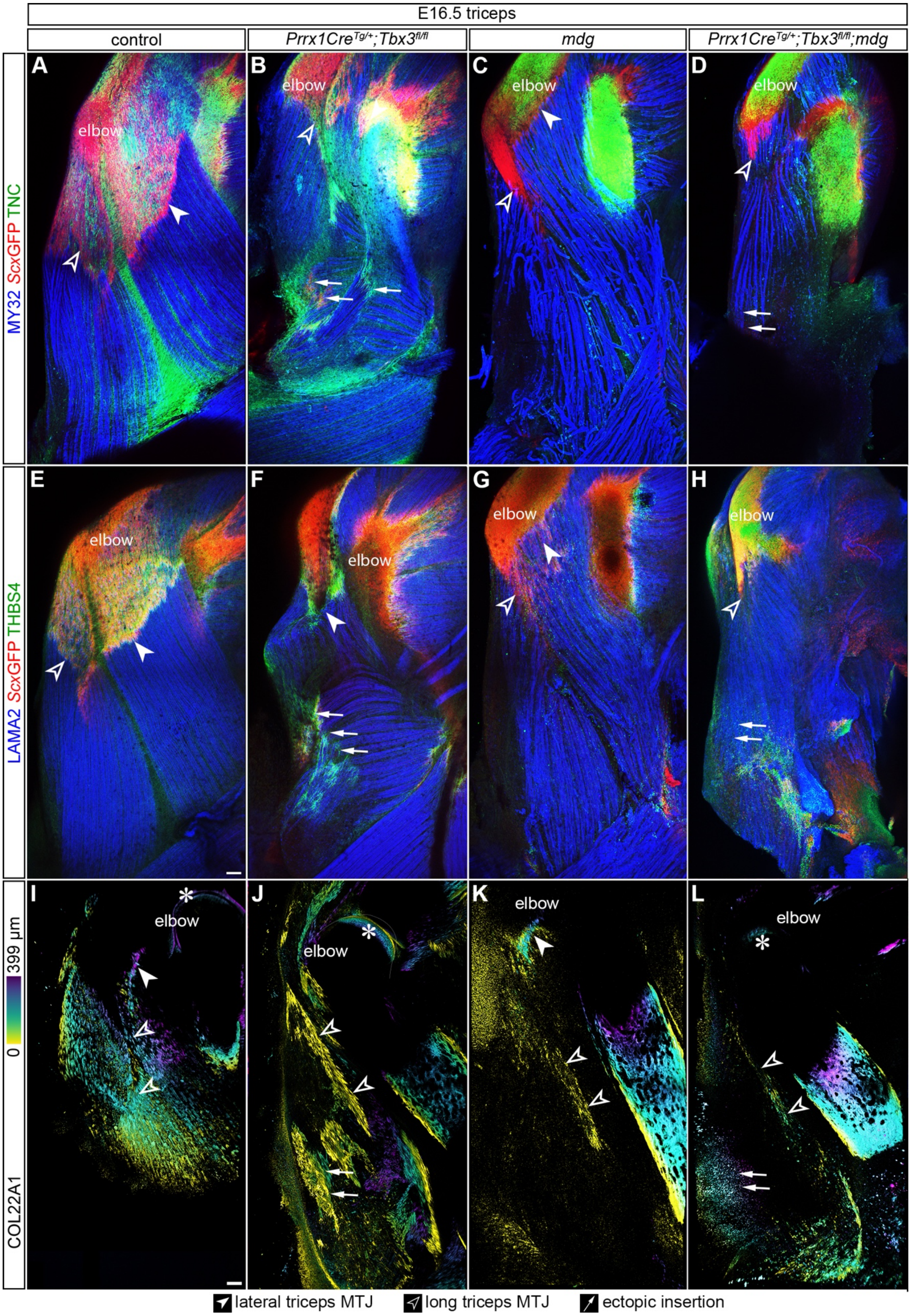
E16.5 triceps
control
*Prrx1Cre*Tg/+*;Tbx3*fl/fl
*mdg*
*Prrx1Cre*Tg/+*;Tbx3*fl/fl*;mdg*
MY32 *Scx*GFP TNC
LAMA2 *Scx*GFP THBS4
COL22A1
0
399 µm
A
B
C
D
E
F
G
H
I
J
K
L
elbow
lateral triceps MTJ
long triceps MTJ
ectopic insertion

**Figure S6: No lateral triceps or COL22A1$^{+}$ MTJ was present in the E16.5 *Prrx1Cre$^{Tg/+}$;Tbx3$^{fl/fl}$;mdg* lateral triceps ectopic insertion.** (**A-H**) Lateral triceps muscle (MY32, LAMA2 = blue) and tendon (TNC, THBS4 = green; *Scx*GFP$^{+}$ = red) formed in the proper location in the control and *mdg* limbs. The lateral triceps was ectopically located in *Prrx1Cre$^{Tg/+}$;Tbx3$^{fl/fl}$* limbs, but was not present in *Prrx1Cre$^{Tg/+}$;Tbx3$^{fl/fl}$;mdg* limbs. (**I-L**) COL22A1$^{+}$ MTJs were present in the long and lateral triceps in both control and *Prrx1Cre$^{Tg/+}$;Tbx3$^{fl/fl}$* limbs at E16.5. COL22A1 was reduced in the lateral and long triceps *mdg* MTJs. In *Prrx1Cre$^{Tg/+}$;Tbx3$^{fl/fl}$;mdg* limbs, COL22A1 was lost at the site of the ectopic insertion, but was present in the long triceps. * Indicates joint surface. A-N: 10× wholemount, z-projection, z = 513 µm (A-D), z = 547 µm (E-G); H-L: 10× decell, 3D rendering, z = 399 µm. Scale bars: 100 µm. Representative images from n = 3 biological replicates.

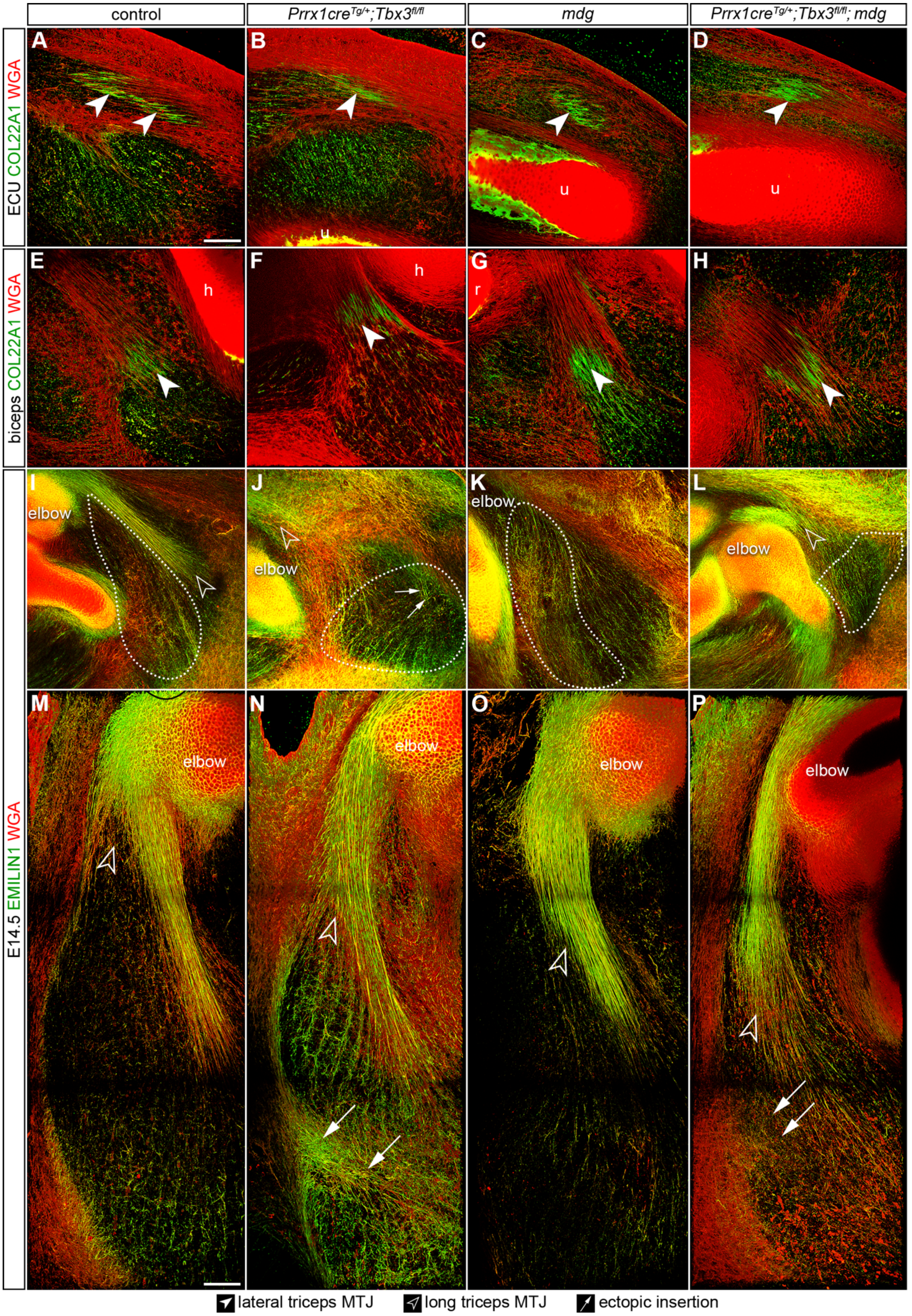

control
Prrx1cre$^{Tg/+}$;Tbx3$^{fl/fl}$
mdg
Prrx1cre$^{Tg/+}$;Tbx3$^{fl/fl}$; mdg
ECU COL22A1 WGA
biceps COL22A1 WGA
E14.5 EMILIN1 WGA
A
B
C
D
E
F
G
H
I
J
K
L
M
N
O
P
u
h
r
elbow
lateral triceps MTJ
long triceps MTJ
ectopic insertion

**Figure S7: The MTJ was not present in the E14.5 *Prrx1Cre$^{Tg/+}$;Tbx3$^{fl/fl}$;mdg* ectopic lateral triceps.** (**A-H**) COL22A1 (green) was expressed in MTJs on the ulnar side of the limb, such as the extensor carpi ulnaris and the bicep brachial (u = ulna, r = radius, h = humerus). (**I-L**) EMILIN1 (green) and WGA (red) was found in the muscle belly fibers of the lateral triceps (dotted line) in control, *Prrx1Cre$^{Tg/+}$;Tbx3$^{fl/fl}$* and *mdg*, but not *Prrx1Cre$^{Tg/+}$;Tbx3$^{fl/fl}$;mdg* limbs. (**M-P**) EMILIN1 was present in control and *Prrx1Cre$^{Tg/+}$;Tbx3$^{fl/fl}$* tendons, whereas it was reduced in *mdg* and absent in the ectopic insertion in the *Prrx1Cre$^{Tg/+}$;Tbx3$^{fl/fl}$;mdg* limbs at E14.5. A-H, M-P: 25× decell, 3D rendering, z = 50 µm (A-D), z = 44 µm (E-H), z = 97 µm (M-P), I-L: 10× decell, z-projection, z = 114 µm. Scale bars: 100 µm. Representative images from n = 3 biological replicates.
